# TCA cycle rewiring underpins implantation and histone acetylation programming

**DOI:** 10.1101/2025.04.25.650409

**Authors:** Eleni Kafkia, David Pladevall-Morera, Lidia Argemi-Muntadas, Gangqi Wang, Roberta Noberini, Sandra Bagés-Arnal, Matthias Anagho-Mattanovich, Rita Silvério-Alves, Tiziana Bonaldi, Ton J. Rabelink, Thomas Moritz, Jan Jakub Żylicz

**Affiliations:** Novo Nordisk Foundation Center for Stem Cell Medicine - reNEW, Department of Biomolecular Sciences, Faculty of Health and Medical Sciences, University of Copenhagen, Copenhagen, Denmark; Novo Nordisk Foundation Center for Basic Metabolic Research, Faculty of Health and Medical Sciences, University of Copenhagen, Copenhagen, Denmark; Department of Internal Medicine (Nephrology) & Einthoven Laboratory of Vascular and Regenerative Medicine, Leiden University Medical Center, Leiden, the Netherlands; Novo Nordisk Foundation Centre for Stem Cell Medicine - reNEW, Leiden University Medical Centre, the Netherlands; Department of Experimental Oncology, IEO, European Institute of Oncology IRCCS, 20139, Milan, Italy; Department of Oncology and Haematology-Oncology, University of Milano, Via Festa del Perdono 7, 20122, Milano, Italy

## Abstract

Metabolism has emerged as a key regulator of stem cell differentiation and their epigenomes. This coupling is particularly evident during the exit from naïve pluripotency in vitro. However, our understanding of the dynamics of the metabolic rewiring especially at implantation remains rudimentary. In this study, we reconstruct the intracellular metabolite routings in pre- and post-implantation mouse embryos and during dynamic pluripotency transitions of cultured stem cells. Our findings reveal that, instead of a simple TCA cycle shutdown, there is a spatio-temporally programmed rewiring of the TCA cycle at implantation. Focusing on the spectrum of pluripotent cells, we identify pyruvate as a key metabolic nexus. Indeed, pyruvate carboxylase and malic enzyme activity establish cyclical carbon flow, which is essential for maintaining a balanced metabolic and transcriptional state and timely exit from naïve pluripotency. Additionally, we discover that formative and primed pluripotent cells exhibit increased glutamine contribution to the TCA cycle, reduced oxidative TCA activity, and reciprocal reductive glutamine metabolism. This metabolic rewiring supports increased histone acetylation turnover, primarily using glutamine as a carbon source, supplemented by pyruvate cycling. Thus, we uncover diverse nutrient strategies that are functionally coupled to epigenome programming and dynamic pluripotency cell state transitions at the time of implantation.

## Introduction

Early mammalian development involves dramatic transcriptional, epigenetic, and metabolic programming, which coordinate successful pregnancy. This is particularly evident when the embryo implants into the uterine lining, and the pluripotent epiblast exits naïve pluripotency to become primed for gastrulation. Unlike transcription and epigenetic regulatory processes, metabolism was traditionally viewed mainly as a housekeeping activity involved in bioenergetic and biosynthetic processes. However, it has become clear that, beyond these vital functions, metabolism also operates as a regulatory pathway influencing cell state and cell fate transitions via its coupling to the epigenome and signaling^1–3^.

In the case of mouse development, fertilization is followed by dynamic rewiring of how the embryo utilizes external nutrients to fuel glycolysis and the tricarboxylic acid (TCA) cycle^4–7^. This process culminates at the blastocyst stage when the embryo exhibits a bimodal metabolic state^6^. Specifically, it uptakes glucose and uses glycolysis intermediates for biosynthetic processes, while funneling pyruvate and lactate into the TCA cycle. Indeed, the blastocyst utilizes oxidative phosphorylation (OxPhos) for energy production, as it shows increased oxygen consumption and upregulates mitochondrial electron transport chain genes^8,9^. How metabolism is wired at later developmental stages remains rather elusive. Transcriptomic analysis suggests a possible downregulation of OxPhos activity and increased reliance on glycolysis^9^. Consistent with this observation, the implanted conceptus exists in a highly oxygen-deprived environment, robustly consuming glucose but little oxygen^8,10^. This is reminiscent of the Warburg effect described previously in multiple cancers^11,12^. Later, at E6.75-7.5 during primitive streak formation, glucose metabolism becomes coupled to somatic lineage allocation^13^. Nevertheless, direct evidence for how metabolism is rewired at implantation is lacking, as is the potential metabolic asymmetry between the lineages of the early embryo. Additionally, it remains unclear how the conceptus can adapt to a low oxygen environment, while maintaining high biosynthetic activity and accurate epigenetic programming.

The progressive development of the epiblast during implantation is accompanied by a dynamic continuum of pluripotent state transitions that are mirrored in vitro by pluripotent stem cell models. The naïve, formative, and primed pluripotency of the pre-, peri-, and post-implantation epiblast can be modeled in vitro using embryonic stem cells (ESC), epiblast-like cells (EpiLC), and epiblast stem cells (EpiSC), respectively^14–17^. By and large, a metabolic switch from bivalent glycolytic and mitochondrial respiration in naïve mouse ESC to exclusively glycolytic in primed mouse EpiSC has been observed based on ECAR measurements in line with in vivo transcriptomic data^18^. On a functional level, α-ketoglutarate (αKG) and glutamine availability are important modulators of naive pluripotency^19–21^. Work with in vitro stem cell models further highlights previously unanticipated complexity and fluidity in how metabolic pathways operate. Indeed, ESCs engage a non-canonical fragmented TCA cycle characterized by increased export of citrate from the mitochondria to the cytoplasm, where it is utilized by the metabolic enzyme ATP-citrate Lyase (ACLY)^22^. While such intricate TCA flows are coupled to cell state transitions in pluripotent stem cells, it remains unclear how metabolism is dynamically rewired. Moreover, it is unclear if in vitro stem cell models recapitulate developmentally-programmed metabolic rewiring, especially since prior studies have been performed under atmospheric oxygen concentration that is manyfold higher than physiological conditions.

Chromatin has emerged as one of the sensors of the intracellular metabolic state thanks to direct coupling between epigenetic modifiers and key central carbon metabolites^1,2^. Implantation constitutes the most dramatic stage of accumulating somatic epigenetic memory in the form of repressive histone and DNA methylation^23–25^. This coincides with rapid enhancer and promoter switching, thus programming a new histone acetylation profile^26^. Depositing such activating chromatin modifications relies on the availability of acetyl-CoA. The levels of this metabolic linchpin fluctuate in different cellular contexts and directly regulate the activity of all histone acetyltransferases (HATs) ^27,28^. Nevertheless, the mechanisms by which pluripotent stem cells sustain adequate bioenergetic and biosynthetic functions under low oxygen, while concurrently maintaining histone acetylation turnover, remain poorly understood.

In this study, we aimed to monitor the dynamic metabolic rewiring that occurs during the exit from naïve pluripotency using mouse E3.5 and E6.5 embryos, as well as in vitro cultured pluripotent stem cell models (**Figure 1A**). We employed ^13^C-labeling techniques under 5% oxygen and utilized recently developed, spatially-resolved matrix-assisted laser desorption/ionization mass spectrometry (MS) imaging (MALDI-MSI) and liquid chromatography MS (LC-MS) as readouts^29–32^. Our findings highlight that, rather than a simple shutdown, there is intricate spatial and time-resolved TCA cycle rewiring when cells exit naïve pluripotency. We identified pyruvate as a key metabolic nexus at the onset of formative pluripotency. Indeed, in addition to the typical pyruvate dehydrogenase entry into the TCA cycle, there is robust pyruvate carboxylase and malic enzyme activity that establish a cyclical carbon flow. Loss-of-function experiments revealed that this pyruvate cycling is crucial for maintaining the metabolic and transcriptional state of ESCs and the robustness of exiting naïve pluripotency. Furthermore, we discovered that formative and primed pluripotent cells establish a metabolic network, characterized by increased glutamine contribution to the TCA cycle and reduced oxidative TCA cycle activity. Pluripotent stem cells also show reversed carbon flow through reductive glutamine metabolism. This carbon routing likely plays a significant role in maintaining epigenome programming, with glutamine, rather than glucose, serving as the most prominent carbon source for the rapid turnover of histone acetylation. Overall, our study reveals unexpected complexity in how central carbon metabolism is rewired at implantation and its dynamic transitions across the spectrum of mouse pluripotency. These diverse nutrient usage strategies are essential for maintaining developmental progression and converge on sustaining rapid epigenome remodeling.

**Figure 1.**
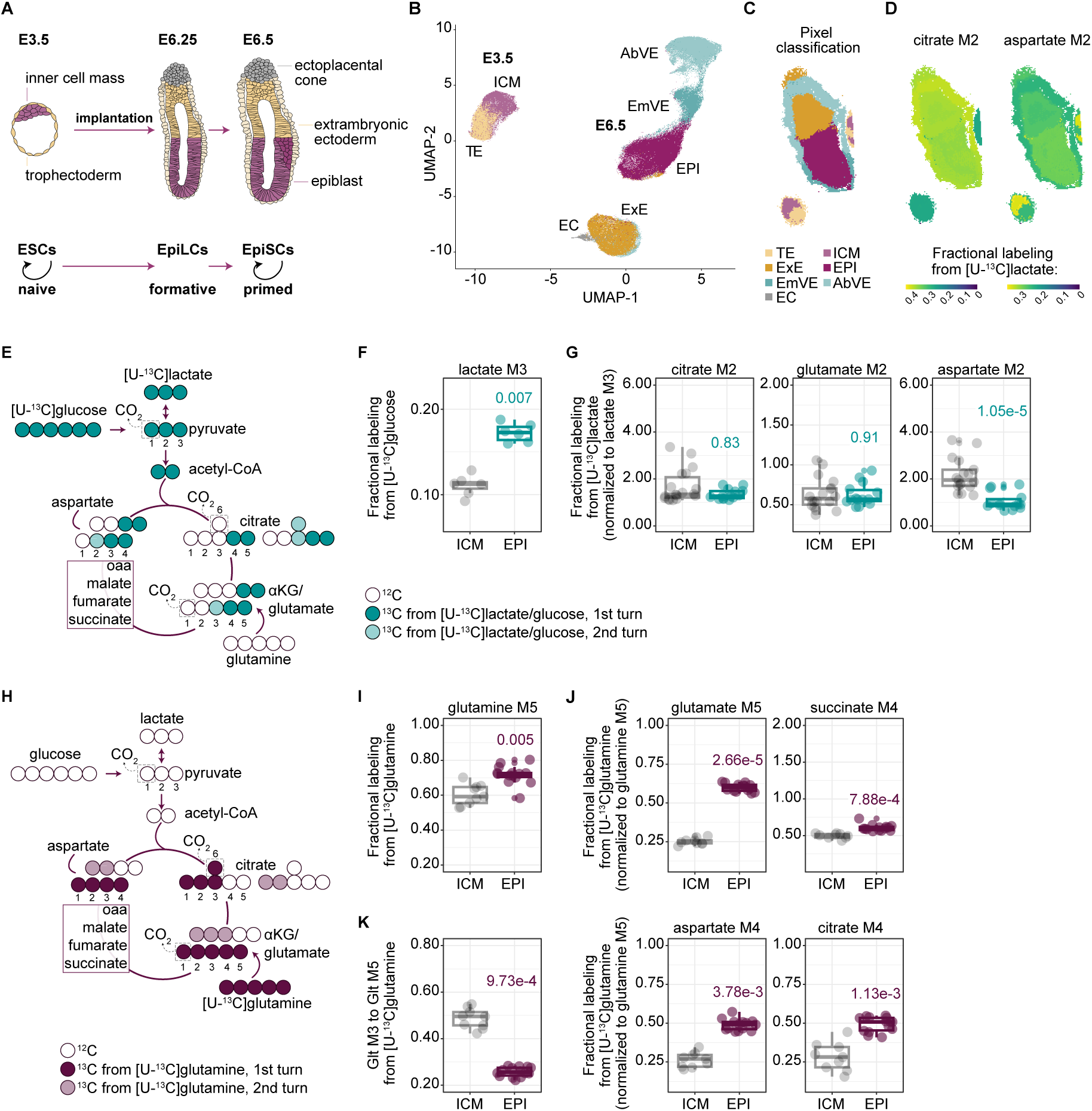
Spatially programmed TCA rewiring underpins embryo implantation. (**A**) Schematic of early mouse embryo development in vivo and corresponding in vitro stem cell models. (**B, C**) UMAP analysis (B) and spatial segmentation (C) of E3.5 and E6.5 embryos based on lipid signals. ICM, inner cell mass; TE, trophectoderm; EPI, epiblast; EmVE, embryonic visceral endoderm; AbVE, abembryonic visceral endoderm; ExE, extraembryonic ectoderm; EC, ectoplacental cone. (**D**) Spatial distribution of M2 citrate and M2 aspartate derived from the [U-^13^C]lactate in E3.5 and E6.5 embryos. (**E**) Schematic illustrating carbon atom transitions in the TCA cycle using [U-^13^C]glucose or [U-^13^C]lactate as tracers. ^13^C carbons are colored. Lighter colored ^13^C carbons derive from the second turn of the TCA cycle. Unlabeled ^12^C carbons are shown as empty. (**F**) Fractional labeling of lactate (M3 isotopologue) derived from [U-^13^C]glucose in the inner cell mass (ICM) and epiblast (EPI) (ICM, n = 7; EPI, n = 6). (**G**) Fractional labeling of the TCA cycle intermediates citrate, glutamate and aspartate derived from [U-^13^C]lactate in ICM and EPI. The M2 isotopologues from the first turn of the TCA cycle are depicted and are normalized to M3 lactate (ICM, n = 18; EPI, n = 14). (**H**) Schematic illustrating carbon atom transitions in the TCA cycle using [U-^13^C]glutamine as a tracer. ^13^C carbons are colored. Lighter colored ^13^C carbons derive from the second turn of the TCA cycle. Unlabeled ^12^C carbons are shown as empty. (**I**) Fractional labeling of glutamine (M5 isotopologue) derived from [U-^13^C]glutamine in ICM and EPI (ICM, n = 9; EPI, n = 14). (**J**) Fractional labeling of the TCA cycle intermediates glutamate, succinate, aspartate and citrate derived from [U-^13^C]glutamine in ICM and EPI. The M4 isotopologues from the first turn of the TCA cycle are depicted and are normalized to M5 glutamine (ICM, n = 9; EPI, n = 14). (**K**) Relative oxidative TCA flux represented as the ratio of M3 glutamate to M5 glutamate from [U-^13^C]glutamine, in ICM and EPI (ICM, n = 9; EPI, n = 14). (**F, G, I-K**) Data points correspond to individual embryos. Statistical significance was assessed using the Kruskal-Wallis test followed by Dunn’s post-hoc test, with p-values adjusted with the Benjamini-Hochberg (BH) correction method.

## Results

### Spatially programmed TCA rewiring underpins embryo implantation

Our current understanding of metabolism in the peri-implantation developmental window is primarily derived from *in vitro* stem cell models and predictions based on *in vivo* transcriptomic data^9,19,21^. These studies have concluded that OxPhos activity decreases in the post-implantation epiblast and primed pluripotent state, and that TCA cycle rewiring is essential for the exit from naïve pluripotency. However, how metabolism rewires in the embryo, especially when the naïve epiblast becomes primed for gastrulation, remains largely unknown. To elucidate this, we adapted a recently developed spatial metabolomics method, which combines MALDI-MSI with stable isotope tracing, for mouse embryos^31,32^. Specifically, we isolated pre-implantation E3.25 blastocysts and post-implantation E6.25 embryos (**Figure 1A**) and cultured them for five hours under 5% O_2_ in chemically-defined media supplemented with uniformly ^13^C labeled nutrients, namely [U-^13^C]glucose, [U-^13^C]lactate, or [U-^13^C]glutamine, which represent key carbon sources for the developing embryo. Afterwards, the samples were flash-frozen and cryosections were used for MALDI-MSI and histology staining. Spatial lipid distributions were used to annotate registered signals to individual lineages and developmental stages. All 5×5 μm pixels were visualized on a uniform manifold approximation and projection (UMAP) plot where they were separated based on lipid signals (**Figure 1B**). Individual Seurat clusters were then annotated based on their stereotypical location (**Figure 1C**). Clusters from the E3.5 blastocysts were annotated as the inner cell mass (ICM) and trophectoderm (TE), whereas E6.5 clusters were annotated as the epiblast (EPI), embryonic visceral endoderm (EmVE), abembryonic VE (AbVE), extraembryonic ectoderm (ExE), and ectoplacental cone (EC) (**Figure 1C**). Next, signals from all pixels originating from the same embryo and the same cluster were averaged and further quantified. Overall, the lipid content efficiently separated known cell types present in the developing embryo (**Figure 1C**).

Using this lipidomic fingerprint, we tracked the incorporation of ^13^C labeled carbon in end glycolytic metabolites and TCA cycle intermediates across distinct lineages (**Figure 1D**). To approximate the labeling of the lower abundance metabolites αKG and oxaloacetate, we used the labeling patterns of glutamate and aspartate, respectively, given their rapid equilibration through transamination reactions. In this context, [U-^13^C]glucose is converted to pyruvate and lactate, both labeled in all three carbons (M3 isotopologues). Pyruvate then enters the TCA cycle through the pyruvate dehydrogenase complex (PDH) yielding acetyl-CoA labeled in two carbons. Subsequently, one round of the TCA cycle will generate metabolites that carry two ^13^C atoms in their carbon backbone (M2 isotopologues) (**Figure 1E**). Tracing [U-^13^C]glucose, we found that E6.5 embryos produce more lactate than E3.5 (**Figure 1F & S1A**), indicating higher glycolytic activity upon implantation. From the [U-^13^C]lactate tracer, we also observed increased lactate uptake at E6.5 compared to E3.5 (**Figure S1B**), and that lactate carbons are further oxidized in the TCA cycle similar to glucose (**Figure S1B**). While citrate, the entry metabolite of the TCA cycle, showed increased labeling at E6.5 relative to E3.5 embryos (**Figure S1B**), this difference was lost after normalizing to lactate labeling, indicating a comparable contribution of lactate to TCA cycle entry between the pre- and post-implantation embryos (**Figure 1G & S2A**). The same pattern was observed for glutamate (**Figure 1G**). In contrast, aspartate, a metabolite further downstream in the TCA cycle indicative of oxaloacetate labeling, showed decreased labeling in E6.5 compared to E3.5 embryos (**Figure 1G & S2A**). This suggested a relative increase in dilution from an unlabeled source downstream of αKG/glutamate, and/or changes in the oxidative TCA cycle in E6.5 embryos compared to E3.5. Interestingly, aspartate M2 labelling differed between E3.5 lineages, with ICM showing a higher percent of labelling compared to TE (**Figure 1D & S1A & S1B**). Together this highlights that there is both spatial as well as temporal control of metabolic flows in the developing mouse embryo.

Beyond glucose and lactate, glutamine is often an important source of TCA cycle carbons. [U-^13^C]glutamine generates glutamate labelled in all five carbons (M5 isotopologues) which then is further deaminated or transaminated to fully labelled αKG. Subsequent oxidation of αKG in the TCA cycle produces metabolites that are labelled in four carbons (M4 isotopologues) by the end of the first round of the cycle (**Figure 1H**). We observed that glutamine uptake was higher at E6.5 compared to E3.5 embryos (**Figure 1I & S1C**). Accordingly, glutamine’s contribution to TCA cycle intermediates (glutamate, succinate, aspartate and citrate M4 isotopologues) was also increased in E6.5 embryos compared to E3.5 (**Figure 1J & S1C & S2B**). Entering the second round of the TCA cycle, M4 citrate loses one ^13^C carbon as CO_2_ forming M3 αKG/glutamate (**Figure 1H**). The ratio of M3 glutamate (second round) to M5 glutamate (first round) was lower in E6.5 embryos compared to E3.5, suggesting reduced oxidative activity in post-implantation embryos (**Figure 1K & S2C**). Within E6.5 embryos, the ectoplacental cone exhibited a higher M3-to-M5 glutamate ratio, and hence oxidative activity, than the epiblast (**Figure S2C**). Additionally, succinate dilution (ratio of M4 succinate to M5 glutamate), showed a trend of being lower in the ICM than in TE, and in ectoplacental cone compared to the epiblast, suggesting reduced contribution of unlabelled sources in these lineages (**Figure S2D**), mirroring the lactate labelling experiment. Taken together, these findings indicate that E6.5 embryos exhibit a more glycolytic metabolism, similar PDH-dependent entry but reduced TCA oxidative activity compared to E3.5 embryos. This metabolic rewiring aligns with the lower oxygen availability in the implanted conceptus^8,10^. While both glucose and glutamine contribute amply to TCA cycle intermediates, glutamine is the major carbon source in the E6.5 embryos. In addition, differential contributions from nutrients other than glucose or glutamine could account for lineage-specific differences in pre- and post-implantation embryos. Overall, our findings reveal that pre-to post-implantation embryonic metabolism does not undergo a simple switch from a bimodal glycolytic and mitochondrial respiration program to an exclusively glycolytic module. Rather, pre- to post-implantation embryo development is marked by intricate TCA cycle rewiring, likely enabling key cellular processes to proceed despite reduced oxygen availability.

### TCA rewiring is a shared metabolic hallmark between embryo’s epiblast and epiblast-like cells

The pre- to post-implantation embryo development is a progressive process that is also tightly linked to a dynamic continuum of pluripotent state transitions, which can be modelled in vitro. To refine our understanding of the sequence of the metabolic transitions occurring *in vivo* during pre- to post-implantation epiblast development in a time-resolved and detailed manner, we next employed a stepwise transition from mouse ESCs to EpiLCs and EpiSCs that marks the embryonic stem cell progression throughout pluripotency. Here, E14 ESCs were cultured in a naive pluripotent state (2i/LIF conditions)^14^. For EpiLC induction, ESCs were plated in N2B27 media that contained Activin A, FGF2 and the canonical WNT signaling inhibitor XAV939 (FAX conditions)^17,33^. The induction to EpiLCs occurred over three days, and the continuous propagation of EpiLCs in FAX conditions resulted in a stable, self-renewing population of EpiSCs (**Figure 2A**)^33^. All the pluripotent stem cells were cultured in the same defined basal N2B27 media and in 5% O_2_ conditions to mimic the physiological *in vivo* low oxygen environment.

**Figure 2.**
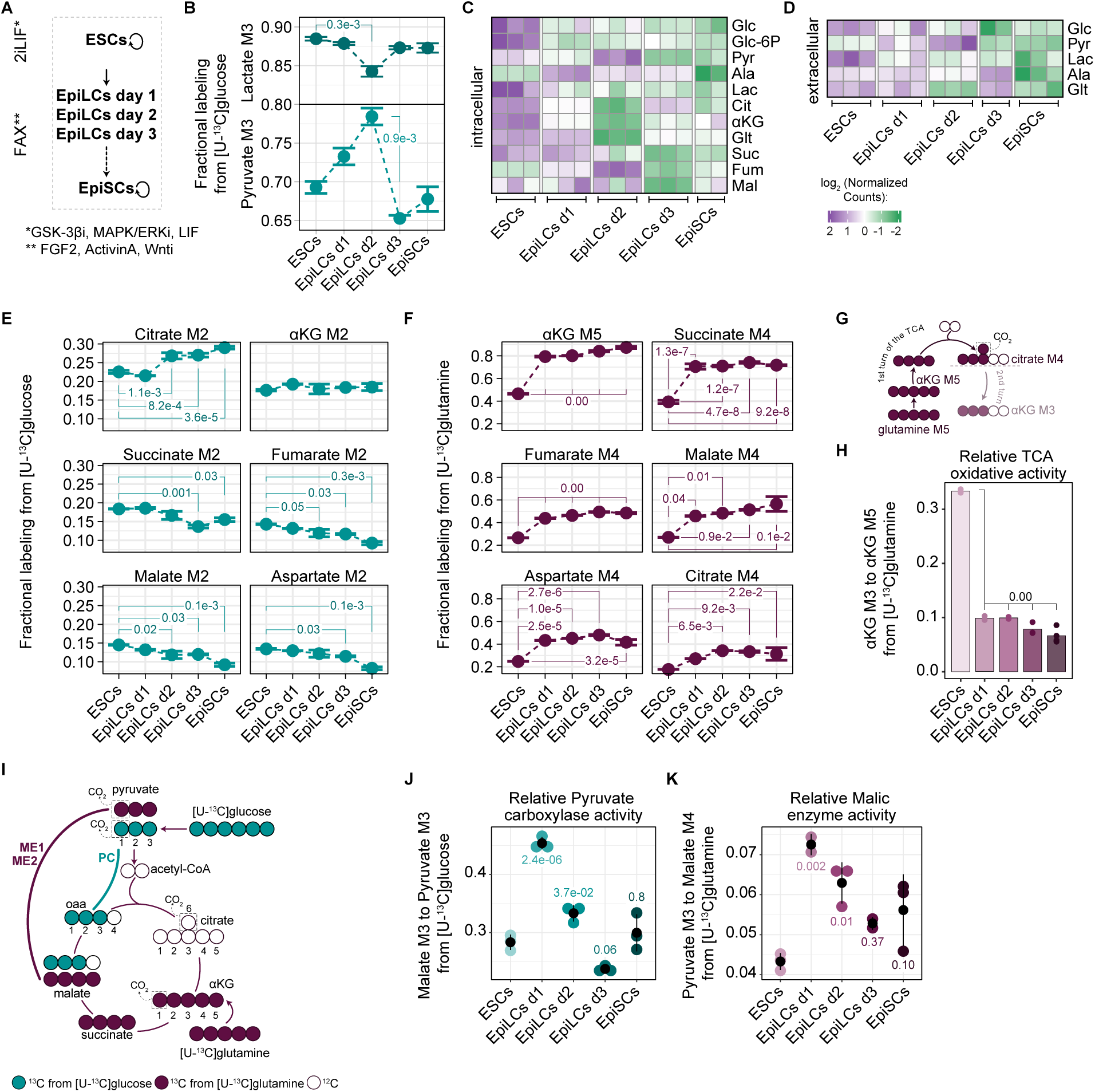
TCA rewiring is a shared metabolic hallmark between embryo’s epiblast and epiblast-like cells. (**A**) Schematic of the *in vitro* stem cell models. GSK-3βi, Glycogen Synthase Kinase-3 beta inhibitor; MAPK/ERKi, Mitogen-Activated Protein Kinases/ Extracellular signal-Regulated Kinases inhibitor; LIF, Leukaemia Inhibitory Factor; FGF2, Fibroblast Growth Factor 2; Wnti, Wnt inhibitor (XAV939). (**B**) Fractional labeling of pyruvate (M3 isotopologue) and lactate (M3 isotopologue) derived from [U- ^13^C]glucose throughout pluripotency progression (n = 3). (**C, D**) Heatmaps of intracellular (C) and extracellular (D) metabolite abundances throughout pluripotency progression. For each metabolite, values represent log2 transformed counts normalized to protein content. Three biological replicates per condition are shown. Glc, glucose; Glc-6P, glucose 6-phosphate; Pyr, pyruvate; Ala, alanine; Lac, lactate; Cit, citrate; αKG, alpha-ketoglutarate; Glt, glutamate; Suc, succinate; Fum, fumarate; Mal, malate. (**E, F**) Fractional labeling of TCA cycle intermediates from [U-^13^C]glucose (E) and [U-^13^C]glutamine (F) throughout pluripotency progression. The depicted isotopologues derive from the first turn of the TCA cycle. E, n = 3; F, n = 3 with the exceptions of glutamate M5 EpiLCs day 1 (n = 2) and malate M4 EpiLCs day 1 and day 2 (n = 2). (**G**) Schematic illustrating carbon atom transitions following the first two turns of the TCA cycle using [U-^13^C]glutamine as a tracer. ^13^C carbons are colored. Lighter colored ^13^C carbons derive from the second turn of the TCA cycle. Unlabeled ^12^C carbons are shown as empty. (**H**) Relative oxidative TCA flux throughout pluripotency progression, represented by the ratio of M3 aKG to M5 aKG from [U-^13^C]glutamine (n = 3). (**I**) Schematic illustrating carbon atom transitions in the TCA cycle following the activities of pyruvate carboxylase (PC) and malic enzymes 1 (ME1) and 2 (ME2). Green and purple indicate ¹³C carbons derived from glucose and glutamine, respectively. Unlabeled ¹²C carbons are shown as empty. (**J**) Relative pyruvate carboxylase activity throughout pluripotency progression, represented by the ratio of M3 malate to M3 pyruvate from [U-^13^C]glucose (n = 3). (**K**) Relative malic enzyme activity throughout pluripotency progression, represented by the ratio of M3 pyruvate to M4 malate from [U-^13^C]glutamine (n = 3). (**B, E, F**) Data represent the mean of three biological replicates ± standard error of the mean (SEM) per condition. (**H**) Data represent the mean of three biological replicates per condition, with individual points corresponding to biological replicates. (**J, K**) Data represent the mean of three biological replicates ± standard deviation (SD) per condition, with colored points indicating individual biological replicates. (**B, E, F, J, K**) Statistical significance was assessed using one-way ANOVA followed by Tukey’s HSD post-hoc test. Levene’s test was used to evaluate homogeneity of variances, and Shapiro-Wilk test was applied to assess the normality of residuals for each metabolite.

To study glucose metabolism, cells were cultivated with [U-^13^C]glucose for five hours, and the ^13^C labeling patterns were analyzed using LC-MS to assess the contribution to metabolic intermediates in glycolysis and the TCA cycle. We observed that lactate labeling (M3 isotopologue) reached above 80% throughout the transition, highlighting the overall highly glycolytic nature of all pluripotent stem cells (**Figure 2B**). However, although EpiSCs have been described as more glycolytic compared to ESCs^18^, M3 lactate remained largely unchanged throughout the transition, except for a transient decrease in EpiLCs on day 2 (**Figure 2B**). Notably, the percentage and kinetics of M3 pyruvate showed a notable discordance with those of M3 lactate (**Figure 2B**). Throughout most of the transition, M3 pyruvate labeling was approximately 20% lower than that of M3 lactate (**Figure 2B**). Furthermore, there was a transient increase in M3 pyruvate, peaking on day 2 of EpiLC induction (**Figure 2B**). This reduction in pyruvate labeling, compared to lactate, suggests that unlabeled carbons from alternative metabolic sources and pathways (e.g., alanine, TCA anaplerosis) converge on pyruvate.

To better understand the glycolytic changes, we used gas chromatography MS (GC-MS) to measure the intracellular and extracellular levels of glucose, pyruvate and lactate, as well as of other central carbon metabolites (**Figure 2C-D**). Throughout the transition, in both intracellular and extracellular pools, lactate decreased, whereas pyruvate fluctuated, with a notable spike on EpiLCs day 2 (**Figure 2C-D**). We also observed a rapid decrease in the levels of the key TCA intermediate αKG upon EpiLC induction, consistent with its function in regulating pluripotency^19,21^. Intracellular and extracellular glucose abundance progressively decreased (**Figure 2C-D**), indicating an increase in glucose consumption during the transition. Despite no significant changes in lactate labeling, the observed increase in glucose uptake suggested that the relative glucose metabolism towards pyruvate and lactate is enhanced during the transition (**Figure S2E, F**). These data align with the embryo labeling results indicating the post-implantation epiblast as more glycolytic. The observed differences in discrete metabolic nodes, such as lactate labeling, between the embryos and pluripotent stem cells likely reflect divergent nutrient availability in both systems.

While the above findings highlight the dynamic changes in glycolytic metabolism during the transition, we next investigated how glucose-derived carbons are incorporated into the TCA cycle. Following the transition, we detected an increase in M2 citrate from EpiLCs day 2 and onwards (**Figure 2E**). The same pattern was observed when the ratio of M2 citrate to M3 pyruvate was calculated to normalize for the upstream changes in glycolysis (**Figure S2G**), suggesting that the induction to EpiLCs is associated with an enhanced pyruvate flux through PDH. Despite this increased glucose entry to the TCA cycle, there was no difference in the abundance of M2 αKG throughout the transition, and the percentage of M2 labeling was consistently lower compared to that of citrate (**Figure 2E**). Metabolites downstream of αKG, namely succinate, fumarate, and malate, showed a progressive loss in M2 isotopologue labelling during the transition (**Figure 2E**). The reduction became even more pronounced at the end of the first round of the TCA cycle, as illustrated by M2 malate and M2 aspartate (**Figure 2E**). This reduced labelling in M2 isotopologues following citrate suggested, at least in part, that a significant proportion of citrate could be utilized in pathways other than the TCA cycle. Consistent with this, it has been demonstrated recently that naïve mouse ESCs possess a fragmented TCA cycle wherein citrate is exported to the cytoplasm for increased utilization by ACLY^22^. Our findings further indicate that this is a feature that characterizes not only the naïve but also the intermediate and primed pluripotent states. These findings are broadly consistent with robust carbon entry into the TCA cycle from glucose/lactate in the E6.5 epiblast and with progressive dilution of labeling at later stages of the TCA cycle.

Several additional mechanisms, and combinations thereof, could further account for the reduced labeling in M2 isotopologues downstream of citrate, including a reduced oxidative TCA activity and the contribution of alternative carbon sources and anaplerotic pathways to TCA intermediates. To further investigate these possibilities, we next used a [U-^13^C]glutamine tracer. We observed significantly higher M4 αKG labeling during the transition, reaching 80% already from EpiLCs day 1 compared to 46% in ESCs (**Figure 2F**). Consistently, the rest of the TCA cycle metabolites exhibited also a significant increase in labelling from glutamine (M4 isotopologues) during the transition compared to ESCs (**Figure 2F**). We also observed that the ratio of M3 αKG (second round) to M5 αKG (first round) was significantly decreased during the transition compared to ESCs (**Figure 2G-H**), indicating a rapid reduction in the oxidative TCA cycle activity. In the context of the primed pluripotent state, these results align with previous studies showing that EpiSCs have reduced mitochondrial respiration and cannot proliferate in the absence of glutamine^18,19^. Together, our data reveal a progressive TCA cycle metabolic rewiring from pre- to post-implantation developmental time wherein formative and primed pluripotency share common metabolic features. More broadly, our findings underscore a fundamental metabolic shift shared between the embryo and stem cells, characterized by robust TCA entry from glucose/lactate, reduced oxidative TCA activity, and enhanced glutamine anaplerosis in the post-implantation stage of development.

### Pyruvate is a distinctive metabolic node at the onset of formative pluripotency

The TCA cycle is supplemented by additional anaplerotic entries and alternative metabolic routes, which replenish key intermediates and provide metabolic robustness under varying physiological conditions. Building on our findings on TCA cycle rewiring, we next sought to determine whether alternative metabolic pathways could further contribute to the observed metabolic shift. Indeed, our analysis revealed a discordant labelling between pyruvate and lactate from the [U-^13^C]glucose tracer (**Figure 2B**), suggesting that carbons stemming from an alternative metabolic route may be converging on pyruvate. Focusing more on the [U-^13^C]glucose tracing experiments, we observed abundant levels of three-carbon labelled (M3 isotopologues) fumarate, malate, aspartate and citrate in ESCs and during the transition (**Figure S2H**). While these isotopologues can be formed from the second round of the TCA cycle and onwards, their labelled fractions were significantly higher relative to that of their precursor metabolites, namely M4 citrate, M3 αKG and M3 succinate (e.g. 19% for M3 malate versus 4.6% for M3 succinate in ESCs) (**Figure S2H**). This finding supported the existence of an alternative glucose entry into the TCA cycle independent of PDH. Pyruvate carboxylase (PC), a key anaplerotic enzyme, converts pyruvate to oxaloacetate (**Figure 2I**), and serves as a primary replenishing route in the liver and some tumors. Our data indicated that substantial glucose-derived ^13^C carbons might enter into the TCA cycle through PC. Notably, this entry significantly spikes on EpiLCs day 1, accounting for as much as 33% of the ^13^C carbon in fumarate, malate and aspartate (**Figure S2H / Table S1**). PC-derived ^13^C carbons eventually flow into citrate, contributing nearly as much as those through the PDH in EpiLCs day 1 (**Figure 1E & S2H**). The same pattern was obtained following normalization of the M3 isotopologues to M3 pyruvate to account for upstream changes in glycolysis (**Figure 2J & S2I-K**).

In addition to PC, malic enzymes (mitochondrial, ME2, and cytoplasmic, ME1) catalyze the conversion of pyruvate to malate, or vice versa as the directionality of the reaction depends on substrate and product concentrations (**Figure 2I**). From the [U-^13^C]glutamine tracer, we detected ^13^C carbons in pyruvate, alanine and lactate, with the highest levels observed in EpiLCs day 1, when normalized to M4 malate (**Figure 2K & S2L, M & Table S1**). The activity of malic enzyme(s) and/or pyruvate carboxylase was further reflected in the presence of M6 citrate in the [U-^13^C]glutamine tracing experiments, whose abundance significantly increased during the transition compared to ESCs (**Figure S2N / Table S1**). Here, glutamine-derived M3 pyruvate produces M2 acetyl-CoA which subsequently condenses with M4 oxaloacetate to generate M6 citrate. Collectively, these findings reveal a substantial glucose contribution to the TCA cycle through an alternative entry possibly involving PC. This anaplerotic influx of pyruvate into the TCA cycle appears to be connected to an efflux of other intermediates from the cycle, such as malate or citrate. Malate can be reconverted to pyruvate through the MEs, completing a pyruvate-malate cycle. An alternate cycle can occur when citrate leaves the mitochondria to be cleaved to acetyl-CoA and oxaloacetate by ACLY. The oxaloacetate can in turn be converted to malate via malate dehydrogenase(s) (MDH), and subsequently back to pyruvate via MEs, forming a pyruvate-citrate cycle. While the relative activity of these two cycles is difficult to estimate the overall pyruvate cycling activity significantly increases at the onset of formative pluripotency and subsequently equalizes to levels similar to those observed in naïve pluripotent state (**Figure 2J, K**).

Although many isotopologues indicative of pyruvate cycling could not be detected in the embryos owing to the limited sensitivity of MALDI-MS, we observed M3 citrate in the [U-^13^C]glucose and [U-^13^C]lactate tracing experiments, in both E3.5 and E6.5 embryos (**Figure S1A, B**). We further detected M5 glutamate using the [U-^13^C]lactate tracer, suggesting a combined contribution from PC and PDH (**Figure 1B**). Taken together, these findings indicated that robust pyruvate re-routing is a defining metabolic feature of stem cells and embryos, positioning pyruvate as a distinctive, developmentally regulated node within central carbon metabolism.

### Pyruvate cycling allows for timely onset of formative pluripotency

The distinct pyruvate re-routing and dynamics prompted us to further investigate its role in pluripotent stem cells and developmental progression. To achieve this, we first sought to validate pyruvate anaplerosis via PC using [1-¹³C]pyruvate, a tracer that enables differentiation between PDH- and PC-dependent glucose entry into the TCA cycle. While PDH activity results in the loss of ^13^C labeled carbon as ^13^CO_2_, PC-mediated entry transfers the ^13^C to oxaloacetate and it is subsequently retained in the TCA intermediates (**Figure 3A**). As such, the presence of M1 malate, fumarate, aspartate and citrate confirmed that pyruvate anaplerosis is robust in stem cells and dynamically regulated during EpiLC induction (**Figure 3B, S4A, B**). Consistent with our findings using the [U-^13^C]glucose tracer, we further observed that EpiLCs day 1 had the highest M1 malate levels (**Figure 3B**), suggesting an increased requirement for pyruvate anaplerosis into the TCA cycle at the onset of formative pluripotency.

**Figure 3.**
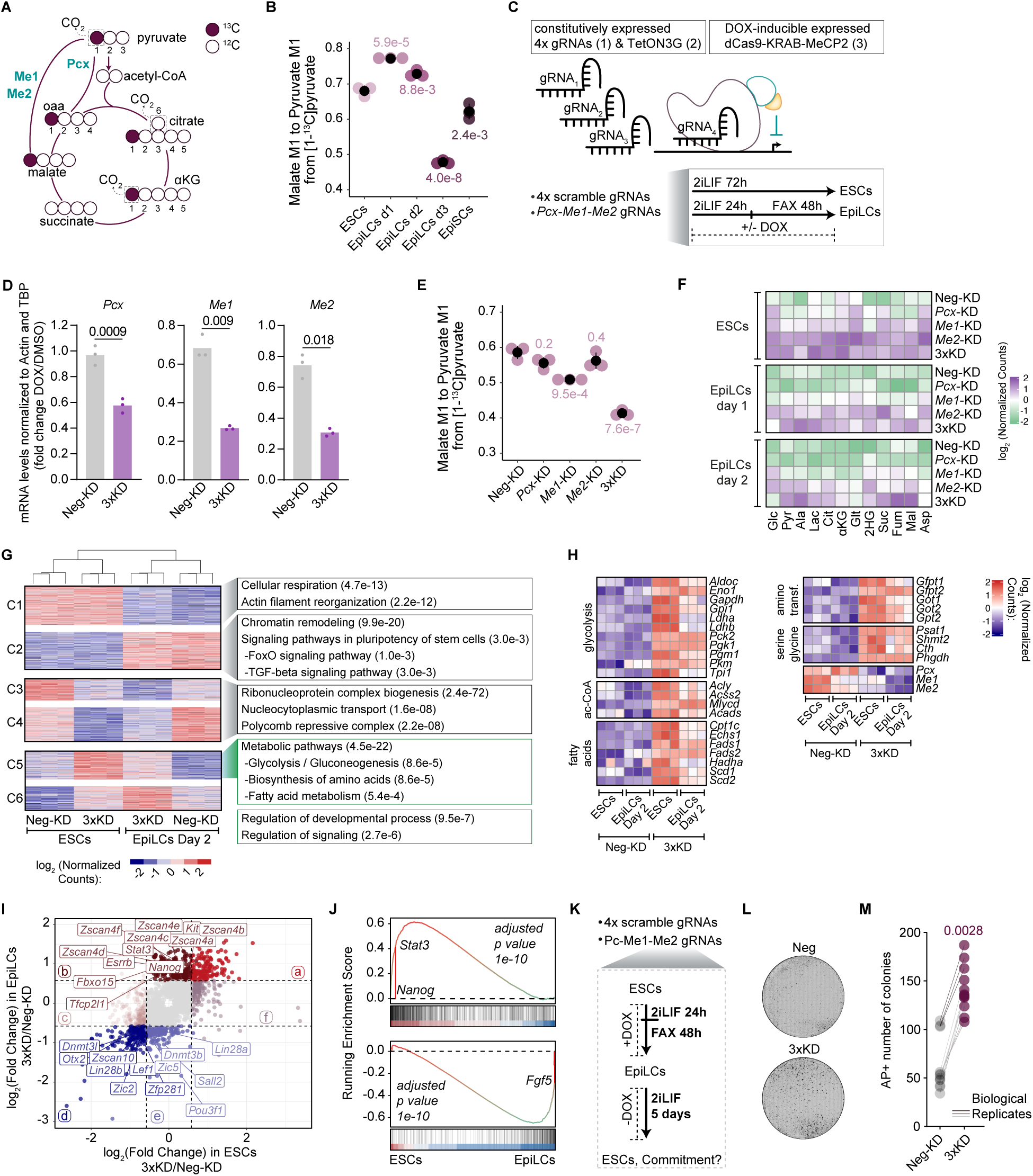
Pyruvate cycling allows for timely onset of formative pluripotency. (**A**) Schematic illustrating carbon atom transitions in TCA cycle using [1-^13^C]pyruvate as a tracer. ^13^C carbons are colored. Unlabeled ¹²C carbons are shown as empty. (**B**) Ratio of M1 malate to M1 pyruvate from [1-^13^C]pyruvate throughout pluripotency progression (n = 3). (**C**) Schematic representation of the E14-DOX-CRISPRi cell line with constitutive TetON3G expression. Upon DOXycycline (DOX) treatment, dCas9-KRAB-MeCP2 is transcribed and guided by the corresponding sgRNAs to bind the transcriptional start site (TSS), silencing target gene(s). Experiments in ESCs were performed in 2iLIF media supplemented with DMSO or DOX for 72 hours. For EpiLCs day 2, cells were cultured for 24 hours in 2iLIF with DMSO or DOX, followed by 48 hours in FAX media with DMSO or DOX. (**D**) RT-qPCR analysis of the indicated genes in scramble control (Neg) and following the repression of all three *Pcx*, *Me1* and *Me2* (3xKD) under DMSO (gray) or DOX (purple) conditions. mRNA levels were normalized to the housekeeping genes *Actin* and *Tbp.* Statistical significance was assessed using two-way ANOVA. Shapiro-Wilk test was used to evaluate residual normality. (**E**) Ratio of M1 malate to M1 pyruvate from [1-^13^C]pyruvate in Neg and following the repression of *Pcx*, *Me1*, *Me2* and all three (3xKD) in EpiLCs day 1 (n = 3). (**F**) Heatmap of intracellular metabolite abundances in Neg and following the repression of *Pcx*, *Me1*, *Me2* and all three (3xKD) in ESCs, EpiLCs day 1 and EpiLCs day 2. For each metabolite, values represent the mean of three biological replicates, with the log2transformed counts normalized to the protein content. Glc, glucose; Glc-6P, glucose 6-phosphate; Pyr, pyruvate; Ala, alanine; Lac, lactate; Cit, citrate; αKG, alpha-ketoglutarate; Glt, glutamate; Suc, succinate; Fum, fumarate; Mal, malate. (**G**) *K-means* clustering of gene expression data in Neg and 3xKD in ESCs and EpiLCs day 2. The most significant gene ontology terms for each cluster are shown (n = 3). (**H**) Expression of metabolic genes in Neg and 3xKD in ESCs and EpiLCs day 2 (n = 3). (**I**) Scatter plot of the log2 transformed fold changes in gene expression in 3xKD compared to Neg in ESCs and EpiLCs day 2. Specific naïve (brown colored) and primed (blue colored) pluripotency genes are highlighted. Dashed lines mark a ± 0.58 log2transformed fold change. a, upregulated in ESCs and EpiLCs in 3xKD compared to Neg; b, upregulated in EpiLCs in 3xKD compared to Neg; c, downregulated in ESCs in 3xKD compared to Neg; d, downregulated in ESCs and EpiLCs in 3xKD compared to Neg; e, downregulated in EpiLCs in 3xKD compared to Neg; f, upregulated in ESCs in 3xKD compared to Neg. (**J**) Gene set enrichment analysis of the significantly upregulated (top) and downregulated (bottom) genes in the 3xKD compared to Neg in EpiLCs day 2. The ranked list was generated from the comparison of Neg ESCs to Neg EpiLCs day 2. (**K**) Experimental setup for assessing the capacity of EpiLCs to revert to ESCs. Neg and 3xKD ESCs were induced to EpiLCs in the presence of DOX. EpiLCs day 2 were plated back in naive (2iLIF) conditions without DOX for 5 days. Subsequently, alkaline phosphatase-positive (AP^+^) colonies were quantified. (**L**) Representative bright-field images of AP^+^ colonies from Neg (top) and 3xKD (bottom) cells. (**M**) Quantification of the number of AP^+^ colonies in the Neg and 3xKD cells following the experimental setup described in (K). (**B, E**) Data represent the mean of three biological replicates ± standard deviation (SD) per condition, with each colored point indicating an individual biological replicate. Statistical significance was assessed using one-way ANOVA followed by Tukey’s HSD post-hoc test. Levene’s test was used to assess variance homogeneity, and the Shapiro-Wilk test was applied to evaluate residual normality. (**M**) Data represent three biological replicates, each with three technical replicates. Statistical significance was determined using an unpaired *t*-test.

To delineate the role that pyruvate routing plays during pluripotency progression, we next aimed to establish a functional framework by developing a multiplex acute loss-of-function system for PC and ME1/ME2 accounting for potential redundancies between these enzymes and cellular adaptations. To this end, we generated an ESC line with a DOXycycline (DOX)-inducible nuclease-dead Cas9 (dCas9) fused to two transcriptional repressor domains: the Krüppel-associated box (KRAB) and the transcription-repression domain of MeCP2, referred to as dCas9-KRAB-MeCP2 (**Figure 3C**). This system has been previously shown to efficiently downregulate genes in HEK-293T and HAP-1 cancer cells^34^. Briefly, in E14 ESCs, the *pTetO-dCas9-KRAB-MeCP2-HaloTag* cassette was knocked into one allele of the safe harbor locus *Tigre* (**Figure S3A**). The other *Tigre* allele was targeted with the *pCAG-TetON3G* constitutive expression cassette. Western blotting, immunofluorescence (IF), and flow cytometry all confirmed robust induction of the dCas9-KRAB-MeCP2-HaloTag 24 hours after DOX addition (**Figure S3B-D**). These cells retained a normal karyotype and were thus tested for knockdown (KD) efficiency (**Figure S3E**). Downregulation of specific genes was achieved by random PiggyBac integration of a sgRNA targeting the transcriptional start site (TSS) of the gene of interest^35^. Scrambled non-targeting sgRNA was used as a negative control throughout the experiments. To validate our new E14-DOX-CRISPRi line, we first integrated three individual sgRNAs targeting the promoter of *Mcm2*, an essential component of the pre-replication complex. Reverse transcription quantitative PCR (RT-qPCR) validated efficient KD, which also led to cell death after 5 days of DOX treatment (**Figure S3F, G**). This E14-DOX-CRISPRi cell line was further functionally validated and was able to generate EpiLCs with similar dynamics to parental E14 cells (**Figure S3H**). Finally, for multiplex knockdowns, we adapted a published sgRNA concatemer vector to the PiggyBac system, allowing for up to 4 unique sgRNAs to be expressed from a single plasmid^36^.

Using the E14-DOX-CRISPRi system, we successfully generated single knockdowns for *Pcx*, *Me1*, and *Me2* (**Figure S3I**). Additionally, with the multiplex system we further established a triple KD (3xKD) cell line by transfecting a vector harboring sgRNAs targeting all three genes (**Figure 3D and S3J**). The CRISPRi cell lines were then induced to EpiLCs day 2, with or without the addition of DOX for 72h, and were supplemented with [1-^13^C]pyruvate for the final five hours. Repression of *Pc* and *Me2* alone caused a slight, albeit not significant, decrease in M1 malate levels (**Figure 3E**). In contrast, knock down of *Me1* and particularly the combined repression of all three enzymes in the 3xKD significantly reduced M1 malate levels, with the latter having the highest impact (**Figure 3E, S4C-D**). These results confirmed that pyruvate rerouting is not only dynamically programmed during the induction to EpiLCs but also mediated through the concerted action of PC and ME1/ME2 enzymes.

To investigate how pyruvate rerouting influences key metabolite levels, we next performed metabolomic analysis in the single KDs and 3xKD in ESCs and through the EpiLC induction (**Figure 3F**, **Table S1**). In all pluripotent states, each KD exerted similar metabolic alterations, albeit with varying degrees of magnitude. Overall, we observed a progressive accumulation of metabolites, with the phenotype being the weakest in *Pcx*-KD, intermediate in *Me1*-KD, and strongest in *Me2*-KD and 3xKD. Compared to negative controls, the most consistent metabolic alteration in the KDs included a significant increase in glucose, glutamate, proline and aspartate abundance, particularly in ESCs. Focusing on metabolites that changed in opposite directions between the different KDs, we found that pyruvate and lactate were increased in all KDs apart from *Pcx*-KD where the levels were significantly reduced. Likewise, downstream of pyruvate, citrate was only increased following *Me1*, *Me2* or 3xKD depletion. Furthermore, we observed that *Pcx* repression resulted in decreased malate levels, an effect that was even more pronounced in EpiLCs day 1, aligning with the higher activity of this pathway at the onset of formative pluripotency. In contrast, depletion of Me1, Me2 and 3xKD led to malate accumulation. This opposite direction on malate abundance reflected the substrate-product dynamics of the underlying metabolic reactions of PC and ME1/ME2, supporting a carbon flow from pyruvate to oxaloacetate/malate via PC and then back to pyruvate via ME1/ME2, consistent with the [U-^13^C]glucose and [U-^13^C]glutamine results. Collectively, these findings further highlight the occurrence of an active pyruvate cycling mediated by the three enzymes. The accumulation of metabolites especially following the repression of all three enzymes suggests compensatory metabolic adaptations and highlights a regulatory role for the pyruvate cycling in bridging the TCA flux with the stem cell demands and balance for carbon, energy, reducing equivalents and overall biosynthetic precursors.

Owing to the high activity of the pyruvate cycling at the onset of formative pluripotency, we next sought to examine its impact on cell state and pluripotency progression by performing RNA sequencing (RNA-seq) in the 3xKD in ESCs and EpiLCs day 2. Unsupervised clustering of differentially expressed genes followed by gene ontology enrichment analysis on each cluster validated that negative control EpiLCs downregulated genes involved in cellular respiration (cluster 1) and upregulated genes linked to chromatin remodeling and signaling in pluripotent stem cells (cluster 2) as expected (**Figure 3G**). Clusters 3 and 4 represented genes that became downregulated in both ESCs and EpiLCs upon repression of *Pcx*, *Me1* and *Me2*. Top enriched GO terms included ribonucleoprotein complex biogenesis and the regulator of developmental genes i.e. Polycomb repressive complex. Clusters 5 and 6 represented genes that were upregulated following the depletion of all three *Pcx*, *Me1* and *Me2* in both ESCs and EpiLCs, and were significantly enriched for metabolic terms (**Figure 3G**). Detailed analysis focusing on central carbon metabolism first confirmed the efficient repression of *Pcx, Me1* and *Me2* in the 3xKD (**Figure 3H**). Additionally, the depletion of all three enzymes was accompanied by robust upregulation of glycolytic genes and glycolytic branching pathways (such as serine/glycine biosynthesis), aminotransferases, fatty acid metabolism, as well as genes involved in acetyl-CoA metabolism such as *Acly* (**Figure 3H**). Amongst the most upregulated glycolytic genes were those directly involved in regulating the pyruvate node and its re-routing. These included pyruvate kinase muscle isoform (*Pkm*), the final rate-limiting glycolytic enzyme that catalyzes the conversion of PEP to pyruvate, and phosphoenolpyruvate carboxykinase 2 (*Pck2*) which also facilitates pyruvate cycling by converting oxaloacetate to PEP. The upregulation of both *Pkm* and *Pck2* was even more pronounced in EpiLCs, and collectively with the overall metabolic gene expression changes suggested a coordinated metabolic adaptation that could enhance glycolytic flux, pyruvate metabolism and reinforce alternative pyruvate cycling routes to compensate for the depletion of this pathway in the 3xKD. These transcriptional changes correlate with the changes observed at the metabolic level in the 3xKD, characterized by the accumulation of glucose and glycolytic intermediates (**Figure 3F**). The transcriptional upregulation of key metabolic genes indicated the involvement of signaling pathways sensing the metabolic state of the cell being modulated by 3xKD. Indeed, there was upregulation of genes associated with signaling in the 3xKD ESCs and EpiLCs (**Figure 3G**). Specifically, gene set enrichment analysis of signaling signatures identified significant deregulation of pathways such as HIF-1, Insulin, and PI3K-Akt (**Figure S3K**). These pathways are known to be sensitive to the metabolic state of the cell and mediate metabolic adaptation^37^.

Furthermore, we focused our analysis on the developmentally-regulated genes altered during the exit from naïve pluripotency. To this end, we plotted the log_2_(fold change) between 3xKD and negative control in ESCs and EpiLCs (**Figure 3I**). This identified a sizeable group of developmental genes specifically misregulated in EpiLCs but not in ESCs (**Figure 3I**). Indeed, we observed an upregulation of naïve pluripotency markers (e.g., *Nanog, Esrrb, Tfcp2l1, Stat3*) and a reciprocal downregulation of primed pluripotency markers (e.g., *Otx2, Dnmt3b, Pou3f1*) in the 3xKD in EpiLCs, indicating a delay in the exit from naïve pluripotency. Gene set enrichment analysis for developmentally regulated genes further supported these findings, confirming that the depletion of pyruvate cycling through PC and ME1/2 disrupts the transition from naive to formative pluripotency (**Figure 3J**). Together, these transcriptomic changes revealed alterations in the metabolic network and developmental program linking pyruvate cycling to the timely onset of formative pluripotency.

To functionally validate the role of PC and ME1/2 in the transition from naive to formative pluripotency, we performed EpiLC reversion assays (**Figure 3K**). Once stem cells have committed to formative or primed pluripotency, they can no longer efficiently revert to naïve pluripotency when placed back into 2i/LIF medium^38,39^. We monitored this by colony formation assay combined with naïve pluripotency-specific staining for alkaline phosphatase activity (AP+). To this end, we dissociated negative control and 3xKD EpiLCs day2 grown in the presence of DOX, plated them in 2i/LIF medium and allowed the colonies to emerge for 5 days in the absence of DOX (**Figure 3K**). As expected, very few cells were able to revert and form AP+ colonies in the negative control cells (**Figure 3L, M**). In contrast, reversion was significantly more efficient upon the depletion of all three enzymes *Pcx* and *Me1/2* (**Figure 3L, M**). Collectively, these findings indicate that the activity of pyruvate cycling is indispensable for a timely commitment to formative pluripotency. Pyruvate cycling serves as a fundamental metabolic circuit by fulfilling the biosynthetic and energy demands of the cells and integrating metabolic homeostasis with developmental timing to safeguard pluripotent transitions.

### Reductive carboxylation of α-ketoglutarate fuels citrate production

Apart from the anaplerotic entry of pyruvate, we have also observed a significant increase in glutamine contribution to the TCA cycle in EpiLCs, EpiSCs, and the E6.5 epiblast compared to naïve pluripotent cells (**Figure 1I, 1J, 2F**). In multiple tumors and the liver, glutamine not only fuels the TCA cycle in the oxidative direction but can also be reductively metabolized^40–44^. In this pathway, isocitrate dehydrogenase carboxylates αKG to form isocitrate, which is then converted to citrate by aconitase (**Figure 4A**). From the [U-^13^C]glutamine tracing experiments, we detected M5 citrate, with labeling reached as high as 30% in EpiLCs day 3 and in EpiSCs (**Figure 4B**), indicating a substantial contribution of reductive glutamine metabolism. Citrate (M5) can be cleaved by ACLY to acetyl-CoA (M2) and oxaloacetate (M3), which subsequently forms malate, fumarate and aspartate (M3) (**Figure 4A**). As such, the presence of M3 malate, fumarate and aspartate further corroborated the occurrence of reductive glutamine carboxylation and activity of ACLY. Indeed, by normalizing M5 citrate to M5 αKG, we quantified relative reductive carboxylation and observed a significant decrease in EpiLCs day 2, along with a trend of increase in EpiSCs (**Figure 4C**). However, the ratio of M3 malate, fumarate, or aspartate to M5 αKG was significantly higher in EpiSCs (**Figure 4D, S4E, F**), suggesting that after αKG is reductively carboxylated to citrate, it is mainly the relative activities of ACLY and/or of the subsequent metabolic enzymes that are higher in EpiSCs compared to ESCs and EpiLCs. To further validate the occurrence of reductive carboxylation, we have performed [1-^13^C]glutamine labelling in ESCs (**Figure 4E**). As expected, we observed M1 labeling at αKG but not at succinate, due to the loss of the M1 label as CO_2_ at the αKG dehydrogenase step. Consistently, we further detected robust M1 labeling at citrate, malate, and fumarate (**Figure 4E**). Taken together, these data indicated a significant presence of reductive carboxylation throughout pluripotency. This metabolic pathway is functionally linked to ACLY activity, which generates acetyl-CoA, a key precursor for fatty acid synthesis and acetyltransferase mediated post-translational modifications. Early development is not the only cellular context in which reductive glutamine metabolism takes place. In certain tissues and cancers, this pathway serves as a major carbon source for lipogenesis, playing an important role in sustaining rapid cell proliferation under hypoxia or conditions of impaired respiration^41–44^. Given their developmental residency in a hypoxic environment and their high proliferative capacity, the pluripotent epiblast may similarly rely on this metabolic pathway.

**Figure 4.**
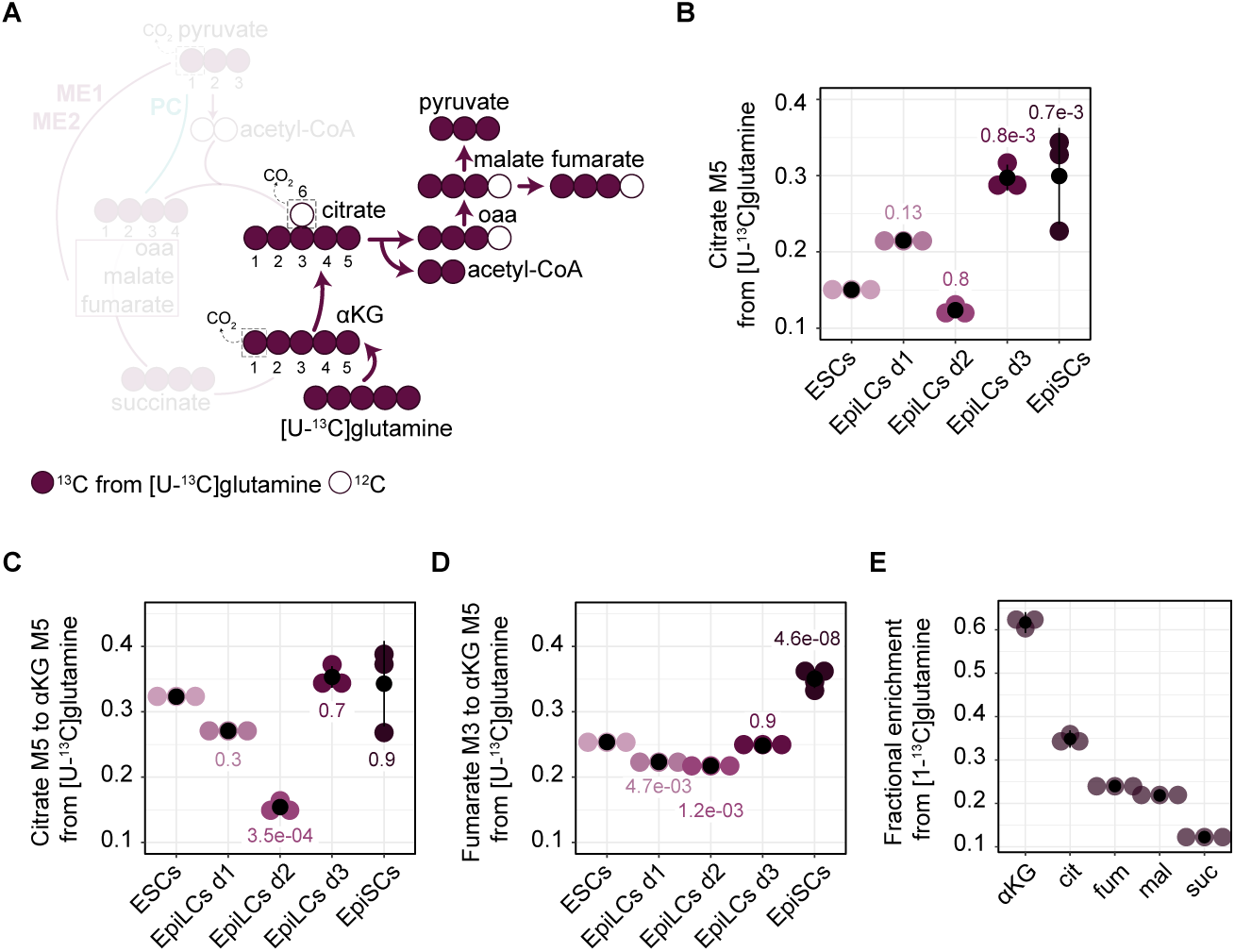
Reductive carboxylation of alpha-ketoglutarate fuels citrate production. **(A)** Schematic illustrating carbon atom transitions during reductive glutamine carboxylation using [U-^13^C]glutamine as a tracer. ^13^C carbons are colored. Unlabeled ¹²C carbons are shown as empty. (**B**) Labeling of citrate (M5 isotopologue) from [U-^13^C]glutamine throughout pluripotency progression (n = 3). (**C**) Relative activity of reductive glutamine carboxylation throughout pluripotency progression, represented by the ratio of M5 citrate to M5 αKG from [U-^13^C]glutamine (n = 3). (**D**) Ratio of M3 fumarate to M5 αKG from [U-^13^C]glutamine throughout pluripotency progression (n = 3). (**E**) Labeling of the TCA cycle intermediates αKG, citrate (cit), fumarate (fum), malate (mal) and succinate (suc) from [1-^13^C]glutamine (n = 3). The main isotopologue, M1, is depicted for each metabolite. (**B-E**) Data represent the mean of three biological replicates ± standard deviation (SD) per condition, with each colored point indicating an individual biological replicate. (**B-D**) Statistical significance was assessed using one-way ANOVA followed by Tukey’s HSD post-hoc test. Levene’s test was used to evaluate homogeneity of variances, and Shapiro-Wilk test was applied to assess the normality of residuals.

### Glutamine is a more prominent source of carbon for histone acetylation than glucose

Owing to the increased glutamine contribution to TCA cycle intermediates in the formative and primed pluripotent states (**Figure 1I, 1J, 2F**), and the robust activity of reductive carboxylation throughout pluripotency (**Figure 4B-E**), we hypothesized that glutamine metabolism through citrate synthesis may contribute substantially to nuclear acetyl-CoA pool used for key regulatory processes, such as histone acetylation. While glucose-derived acetyl-CoA has traditionally been considered the primary source for histone acetylation^45,46^, we sought to evaluate the relative contributions of glutamine and glucose to this process to test our hypothesis. Indeed, from the ^13^C tracing metabolomics experiments (**Figure 2E & 2F**), we calculated that the total glutamine contribution to citrate surpassed that of glucose from the onset of formative pluripotency (**Figure 5A, B**). To quantify this effect directly at the histone acetylation level, we next cultured ESCs, EpiLCs (day 2) and EpiSCs with [U-^13^C]glucose or [U-^13^C]glutamine for 5 hours and analyzed the ^13^C incorporation in acetylated histone peptides by mass spectrometry (**Figure 5C & 5D**). Since the carbon for histone methylation originates mainly from extracellular methionine, we did not monitor ^13^C labelling into the methyl groups. As such, this approach enabled us to determine both the relative abundance of individual histone post-translational modifications and the relative contribution of glucose and glutamine to acetylation (**Figure 5C, D, S4G**).

**Figure 5.**
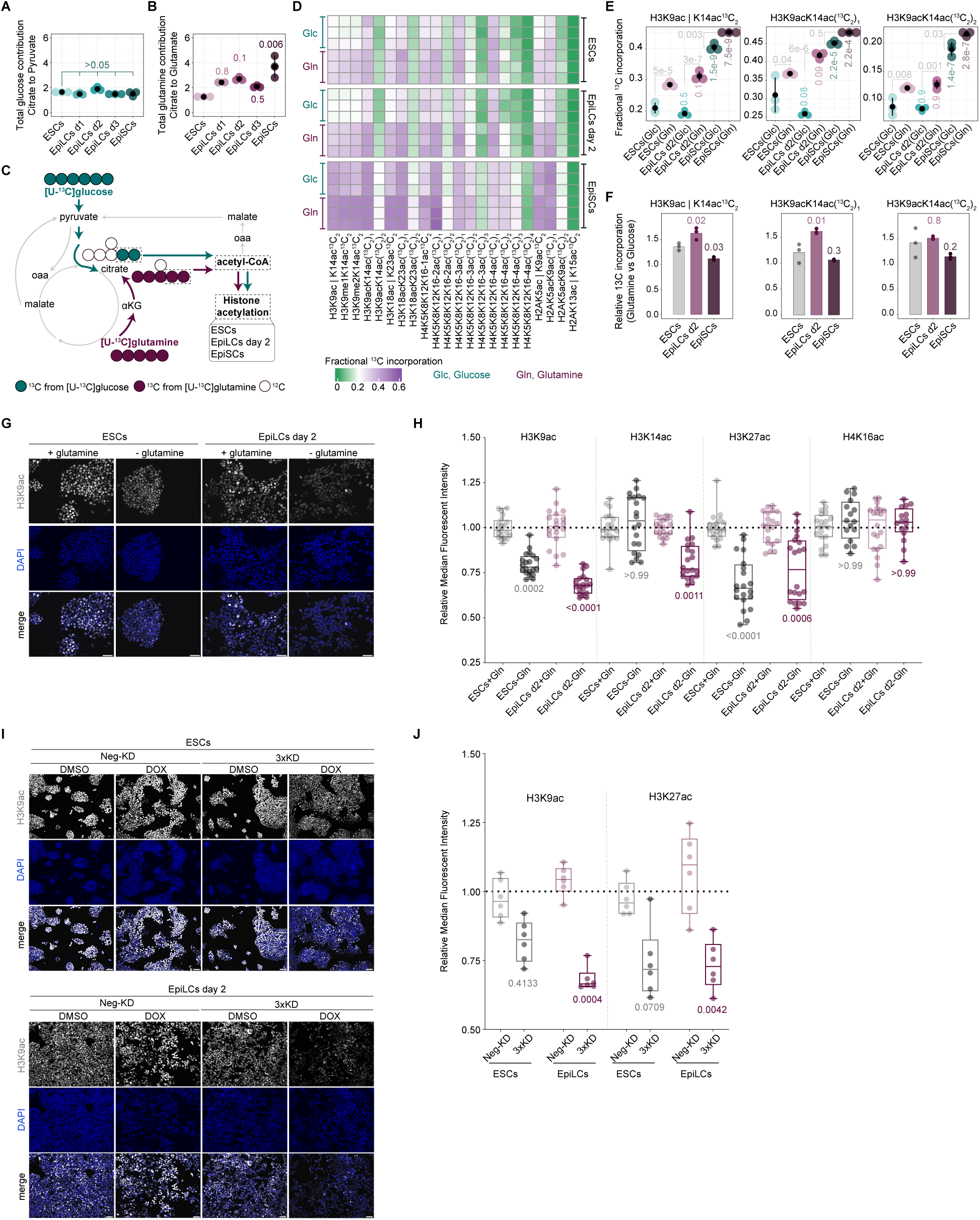
Glutamine is a more prominent source of carbon for histone acetylation than glucose. (**A, B**) Total carbon contribution of glucose (A) and glutamine (B) to citrate throughout pluripotency. The values have been normalized to the total carbon contribution of glucose to pyruvate (A), and to the total carbon contribution of glutamine to glutamate (B). The carbon contributions are based on [U-^13^C]glucose and [U-^13^C]glutamine tracing (n = 3). (**C**) Schematic illustrating carbon atom labeling in citrate using [U-^13^C]glucose or [U-^13^C]glutamine. Citrate is converted to acetyl-CoA which subsequently can be utilized for histone acetylation. To detect the incorporation of glucose or glutamine carbons to histone acetylated peptides, ESCs, EpiLCs day 2 and EpiSCs were cultured for 5 hours in the presence of either [U-^13^C] glucose or [U-^13^C] glutamine prior to downstream sample processing and analysis of histone acetylation by MS. Green and purple indicate ¹³C carbons derived from glucose and glutamine, respectively. Unlabeled ¹²C carbons are shown as empty. (**D**) Heatmap showing the fractional ^13^C-incorporation for differentially modified histone peptides derived from [U-^13^C] glucose (Glc) or [U-^13^C] glutamine (Gln) in ESCs, EpiLCs day 2 and EpiSCs. Residues separated by “l” indicate the measurement of one residue or the other (n = 3). (**E**) %^13^C-incorporation for the indicated histone post-translational modification residues derived from [U-^13^C] glucose or [U-^13^C] glutamine in ESCs, EpiLCs day 2 and EpiSCs (n = 3). (**F**) Ratio of the glutamine to glucose %^13^C-incorporation for the indicated histone post-translational modifications in ESCs, EpiLCs day 2 and EpiSCs (n = 3). (**G**) Representative IF images of H3K9ac (gray) in ESCs and EpiLCs day 2 under control conditions or following 5h of glutamine depletion. Scale bar = 50µm. (**H**) Relative median fluorescent intensity of indicated histone acetylation marks in ESCs and EpiLCs day 2 under control conditions or following 5h of glutamine depletion. For each acetylation mark, the median fluorescent intensity for each data point was normalized to the average median fluorescent intensity of the corresponding control. Data corresponds to 5 biological replicates with each biological replicate including 4 technical replicates (n>20.000 cells). (**I**) Representative IF images of H3K9ac (gray) in Neg and 3xKD cell lines in ESCs and EpiLCs day 2 in the presence of DMSO or DOX for 72 hours, grown in the corresponding cell culture media. Scale bar = 50µm. (**J**) Relative median fluorescent intensity of indicated histone acetylation marks in Neg and 3xKD cell lines in ESCs and EpiLCs day 2. For each histone acetylation mark, the fluorescent intensity was calculated by measuring the median intensity of DOX-treated cells normalized to their DMSO counterparts. Data corresponds to 6 biological replicates (n>20.000 cells). (**A, B, E, F**) Data represent the mean of three biological replicates (A, B, E, F) ± standard deviation (SD) (A, B, E) per condition, with each colored point indicating an individual biological replicate. (**A, B, E, F**) Statistical significance was assessed using one-way ANOVA (A, B, F) or two-way ANOVA (E) followed by Tukey’s HSD post-hoc test. Levene’s test was used to evaluate homogeneity of variances, and Shapiro-Wilk test was applied to assess the normality of residuals. (**H, J**) Median, maximum and minimum values are indicated. Statistical significance was assessed by Kruskal-Wallis test followed by Dunn’s post-hoc test, with p-values adjusted with the Benjamini-Hochberg (BH) correction method.

By quantifying the relative abundance of different histone modifications, we first confirmed a robust increase in H3K9me2 and a loss of H3K27me3 at the onset of formative pluripotency (**Figure S4G**), consistent with previous reports^24^. Additionally, we observed that global levels of multiple histone acetylation marks are dynamic, with H3K9ac and H3K14ac decreasing in EpiSCs compared to ESCs. These findings aligned with key epigenetic programming events during implantation and further highlighted the extensive reorganization underlying the global histone acetylome^26^. Next, to elucidate the metabolic source of histone acetylation, we analyzed the ^13^C incorporation in acetylated histone peptides and observed that across all pluripotent states, glutamine-derived carbon surpassed the glucose-derived carbon (**Figure 5D & S4H**). This ^13^C incorporation progressively increased from the naïve to the formative and the primed pluripotent state (**Figure 5D, E & S4H**). Specifically, when comparing ESCs to EpiSCs, the ^13^C labeling of the mono-acetylated H3K9K14 peptide (H3K9ac|K14ac) increased from 28.4% to 45.3% from the [U-^13^C]glutamine tracer, and from 20.1% to 40.0% from the [U-^13^C]glucose tracer (**Figure 5E**). Similar trends were observed across all peptides, albeit with varying levels of labeling. Thus, there is more robust usage of carbon from glutamine to acetylate histones compared to glucose throughout pluripotency. Moreover, there was an increased overall histone acetylation turnover in EpiSCs compared to ESCs and EpiLCs, as demonstrated by the high levels of labeling. Indeed, in EpiSC for multiple peptides, the combined contribution of glucose- and glutamine-derived carbons to acetyl groups exceeds 80% (e.g. H3K9ac|K14ac: 85.7%; H3K18ac|K23ac: 82.4%). This indicated that there are no alternative major sources of acetyl-CoA used by histone acetyltransferases besides glutamine and glucose. Furthermore, for marks such as H3K9ac and H3K14ac, the nearly complete turnover within the five-hour labeling window highlighted the rapid histone deacetylation and reacetylation dynamics. Finally, we assessed whether the ^13^C incorporation from glutamine to histone acetylation changed relative to glucose. Indeed, there was a modest but significant increase in the glutamine to glucose 13C incorporation ratio in EpiLCs day 2 (**Figure 5F**). The overall trend suggests that increased glutamine entry into the TCA cycle is balanced by the increased pyruvate entry at the onset of formative pluripotency (**Figure 2E**). Together, these findings reveal that glutamine contribution to the histone acetylation backbone exceeds that of glucose.

To further validate this, we deprived ESCs and EpiLCs (day 2) of glutamine for five hours and assessed the levels of specific histone acetylation marks by IF. In line with the previous observations, the relative fluorescence intensity of key histone acetylation marks (H3K9ac, H3K14ac, and H3K27ac) was significantly reduced in glutamine-deprived EpiLCs compared to replete conditions (**Figure 5G-H**). Similarly, glutamine-deprived ESCs exhibited a significant decrease in H3K9ac and H3K27ac relative to their replete counterparts (**Figure 5G-H**). In contrast, H4K16ac remained unchanged across all conditions (**Figure 5H**). Although H4K16ac was not uniquely identifiable by MS, these findings suggest that it likely exhibits a lower turnover compared to the other examined acetylation marks. Together, these results demonstrated that glutamine depletion leads to a significant reduction in histone acetylation marks, reinforcing its essential role in sustaining the dynamic histone acetylation turnover associated with pluripotency transitions.

Given the increased contribution of both glutamine- and glucose-derived carbons to histone acetylation, we hypothesized that this process could be metabolically coupled to pyruvate cycling, which replenishes TCA cycle intermediates, thereby supporting acetyl-CoA availability to histones. To investigate this, we utilized the CRISPRi 3xKD cell line to deplete all three *Pcx*, *Me1* and *Me2* in ESCs and EpiLCs (day 2) and quantified H3K9ac and H3K27ac abundance by IF. Indeed, we observed that depletion of all three enzymes led to a significant reduction in these histone acetylation marks (**Figure 5I-J**), thus confirming that pyruvate cycling, along with reductive glutamine metabolism, is essential for histone acetylation. Collectively, these findings suggested that pyruvate cycling maintains a steady supply of oxaloacetate and citrate, optimizing TCA flux and enhancing acetyl-CoA synthesis, which in turn supports histone acetylation dynamics in naïve and formative pluripotency.

## Discussion

Our study reveals that central carbon metabolism undergoes a dynamic rewiring at the time of implantation. By using spatially resolved ^13^C isotope tracing in pre- and post-implantation mouse embryos, alongside ^13^C isotope tracing in *in vitro* stem cells spanning the entire pluripotency spectrum, we delineate both temporal and spatial metabolic circuitries that underpin early embryo development. Previous studies, based on *in vivo* transcriptomics data and ECAR measurements in naïve ESCs and primed EpiSCs, suggested a metabolic switch from a dual glycolytic-oxidative metabolism in the pre-implantation stage to an exclusively glycolytic one post-implantation^9,18^. Here we uncover that, under physiological oxygen conditions, rather than a simple metabolic switch, there is an intricate rewiring of the TCA cycle. Initially, in the blastocyst, there is substantial glucose flux into the TCA cycle, with lineage-specific differences between the ICM and TE. In line with this, glucose begins contributing to the TCA intermediates only at the blastocyst stage and not earlier in development^6^. As implantation commences and cells transition out of the naïve state, we observe a reduction in oxidative TCA activity, which is consistent with the hypoxic *in utero* environment. However, notably, there is no overall shutdown of the TCA cycle. Rather, there is an increase in glucose consumption accompanied by increased entry into the TCA cycle through PDH. This high glucose TCA entry likely supports a biosynthetic role, as citrate can be exported to the cytosol, where it is cleaved by ACLY to produce acetyl-CoA^46^. Indeed, *Acly* knockout ESCs fail to efficiently exit naïve pluripotency upon 2iLIF withdrawal^22^. This highlights that, at the time of implantation, when pluripotent cells accelerate their cellular divisions, they likely require a robust source of acetyl-CoA for lipid biosynthesis and protein post-translational modifications.

In addition to glucose TCA entry through PDH, our data reveal a significant alternative route through PC. We observe PC activity in both pre- and post-implantation embryos and throughout the pluripotency continuum. Consistent with our findings, ESCs cultured under atmospheric oxygen levels also show some PC activity, albeit at manyfold lower levels^20^. Additionally, during pre-implantation development, PC activity increases at the blastocyst stage^6^. Notably, we find that anaplerotic flux through PC increases significantly at the onset of formative pluripotency, suggesting that anaplerotic entry of pyruvate into the TCA cycle could be developmentally regulated. The importance of PC in cellular physiology has been mainly described beyond embryonic contexts. In the liver, PC is essential in maintaining TCA cycle activity by replenishing metabolic intermediates, in gluconeogenesis and in redox homeostasis^47^. In certain tumors *in vivo*, PC has emerged as the primary TCA anaplerotic route enabling glutamine-independent growth, with its deletion compromising tumor formation^48–52^. Broadly, PC supports biosynthesis and thus metabolic flexibility, processes that are critical not only for tumorigenesis and hepatic metabolism, but also, as our findings suggest, for stem cell state transitions. Notably, we find that this anaplerotic flux through PC is also accompanied by a reverse carbon flow via malic enzymes which convert malate back into pyruvate. Similar to PC, malic enzyme activity spikes at the onset of formative pluripotency. This coordinated activity constitutes a pyruvate cycling pathway, in which malate is recycled into pyruvate and re-enters the TCA cycle via both PDH and PC. Interestingly, a similar metabolic cycle has been described in normoxic differentiating adipocytes where it has been suggested to serve NADPH production and cytosolic acetyl-CoA biosynthesis for lipids^53,54^. Functionally, we demonstrate that disrupting pyruvate cycling through simultaneous knockdown of *Pcx*, *Me1*, and *Me2* impairs the rate of transition from naïve to formative pluripotency, highlighting metabolic coupling to cell state transitions. Mechanistically, the reasons for this could be manifold. On one hand, we find that in ESCs devoid of pyruvate cycling there is an accumulation of αKG, a metabolite known to sustain naïve pluripotency^19,21^. In a manner analogous to adipocytes, pyruvate cycling could support the biosynthetic demands by coupling energy and acetyl-CoA availability through ACLY, whose activity is also important during the exit from naïve pluripotency^22^. Finally, repression of pyruvate cycling results in an altered signaling milieu which could stabilize naivety. Therefore, pyruvate through its cycling emerges as a crucial metabolic node during the exit from naivety, likely by integrating stem cell demands for carbon and energy and being tightly coupled to cell state transitions.

Upstream of pyruvate, differential glucose metabolism has been shown to regulate cell fate decisions during gastrulation^13^. At this developmental stage, hexosamine biosynthetic pathway (HBP) activity is essential for primitive streak elongation, whereas inhibition of the last glycolytic step, namely the phosphoenolpyruvate to pyruvate conversion by pyruvate kinase M2 (PKM2), has minimal impact. Notably, our findings reveal that the post-implantation epiblast can utilize exogenous lactate, which can be converted to pyruvate and subsequently fuel the TCA cycle. This route bypasses the PKM2-mediated glycolytic flux, raising thus the possibility that pyruvate metabolism may play a broader developmental role, beyond its involvement in naïve pluripotency, potentially contributing to later stages of development, including gastrulation.

Beyond pyruvate metabolism, our findings further show that post-implantation embryo development and the transition to formative and primed pluripotent states are characterized by an increase in glutamine anaplerosis into the TCA cycle. This is consistent with prior observations that Serum/LIF cultured ESCs, as well as EpiSCs, cannot proliferate in the absence of glutamine ^19,20^. Most notably, our data also points towards a substantial glutamine reductive carboxylation present throughout the pluripotency continuum. This alternative metabolic route, where glutamine-derived αKG is reductively carboxylated to isocitrate and citrate, is often coupled to ACLY activity for the biosynthesis of acetyl-CoA which subsequently can be used for lipids. The reductive glutamine carboxylation has been well documented in highly proliferative systems such as cancers, where it supports biosynthetic purposes, particularly under hypoxic conditions or when mitochondrial respiration is impaired^41–44^. In the context of stem cells, a low level of reductive glutamine metabolism has been observed ^55,56^. However, our data demonstrate that under physiological oxygen conditions reductive glutamine metabolism is manyfold more prominent. Therefore, we find that reductive glutamine metabolism is a core feature of pluripotent metabolism which, in a manner analogous to cancer, could be employed as an efficient biosynthetic strategy providing acetyl-CoA for lipid and/or acetylation-dependent processes.

Overall, our ^13^C labeling, loss-of-function, and glutamine starvation experiments reveal that pyruvate and glutamine TCA anaplerosis, coupled with reductive carboxylation, sustain the nuclear pool of acetyl-CoA. Indeed, only the acetyl-CoA in the nuclear compartment can robustly contribute to histone acetylation, which is necessary to sustain transcription. However, it remains unclear to what extent it is produced locally, or it diffuses from the cytoplasm. Nuclear acetyl-CoA can originate from pyruvate, citrate, acetate, or lipid metabolism^57^. In most cells, including pluripotent stem cells, citrate is the major source of nuclear acetyl-CoA used for histone acetylation^46,58^. Indeed, in HEK293T cells, acetate does not robustly contribute to histone acetylation; rather, it is almost entirely dependent on glucose-derived citrate^45^. In human primed pluripotent stem cells, citrate is also the major source of histone acetylation, but its depletion can be compensated for by extracellular acetate supplementation^58^. Here, we demonstrate that in all tested mouse pluripotent stem cell models, glutamine reductive metabolism is the major source of carbon for histone acetylation, surpassing that derived from glycolysis, which is not typically observed in HEK293T cells^45^. There is still sparse direct evidence for glutamine being a major source of carbon for histone acetylation in other systems. However, some loss-of-function experiments have found that disrupting glutamine metabolism in chondrocytes diminishes their histone acetylation levels^59^. Our results indicate that at a crucial developmental junction, glutamine is the major source of carbon for histone acetylation. In addition, our data also show that pyruvate cycling is vital for sustaining histone acetylation, especially in formative pluripotency. These results align with findings from the liver and cancer, where PC activity is crucial for acetyl-CoA production and fatty acid synthesis. Thus, dynamic TCA anaplerosis converges at the level of nuclear and cytoplasmic acetyl-CoA, in part used to fuel histone acetylation.

Our histone PTM ^13^C labeling uncovered a previously unanticipated acceleration in histone acetylation turnover upon exit from naïve pluripotency. In agreement with previous work, we identify H3K9ac and H3K14ac as some of the most dynamic marks^60^. Increased turnover of histone acetylation might indicate a general increase in histone turnover in EpiSCs or alternatively stimulated histone deacetylase activity. The former hypothesis is supported by an accelerated cell cycle in the peri-gastrulation embryo^61^. The alternative is supported by the overall decrease in histone acetylation levels in EpiSCs (**Figure S4G**), coinciding with dramatic histone acetylation repositioning during the ESC-to-EpiLC transition^26^. Along this line, the active enhancer mark H3K27ac is removed from ESC-specific genes and deposited on EpiLC enhancers during the onset of formative pluripotency^26,62^. We show that lack of functional pyruvate cycling by loss of *Pcx/Me1/Me2* leads to decrease levels of both H3K9ac and H3K27ac in naïve and formative stem cells and simultaneously impacts the exit from naïve pluripotency. Indeed, primed and formative cells need to rapidly respond to differentiation cues, a process possibly enhanced by high histone acetylation turnover mediated by histone deacetylases^63,64^.

Overall, our study reveals unexpected complexity in how central carbon metabolism is rewired at implantation. These diverse nutrient usage strategies are essential for maintaining developmental progression and epigenome remodeling. Together this highlights similarities in how the embryo adapts to environmental changes at implantation and how multiple cancers may exploit these pathways to rewire their metabolism in a hostile niche.

## Resource availability Lead contact

Further information and requests for resources and reagents should be directed to and will be fulfilled by the lead contact, Jan Jakub Żylicz.

## Data and code availability

Sequencing data that support the findings of this study have been deposited in the Gene Expression Omnibus (GEO) under accession code GSE294457.

## Acknowledgments

This work was supported by grants from Novo Nordisk Fonden (NNF)(NNF21CC0073729, NNF23SA0084103), Lundbeckfonden (Lundbeck Foundation) (R345-2020-1497, R380-2021-1519), European Research Council(ERC)(101077271) and Danmarks Frie Forskningsfond (DFF) (0169-00031B). We are grateful to Robert Bone for critical feedback and to Alessandro Ghiringhelli, Viktoria Lavro for experimental support. We thank Anne Wenzel for computational support. We thank reNEW and CPR platforms for technical expertise, support and use of equipment in particular: H. Wollmann, M. Michaut, J. Bulkescher, G. Dela Cruz and A. Kalvisa.

## Author contributions

Conceptualization, EK, DP-M, JJZ

Methodology, EK, DP-M, GW, RN, TB, TR, TM, JJZ

Investigation, EK, DP-M, LA-M, GW, RN, SB-A, MA-M, RA-S, JJZ

Formal analysis, EK, DP-M, LA-M, GW, RN, MA-M, TM, JJZ

Data Curation, EK, GW, RN, TM

Visualization, EK, DP-M, GW, RN, JJZ

Writing – original draft preparation, EK, DP-M, JJZ

Writing – review and editing, EK, DP-M, LA-M, GW, RN, RA-S, TB, TR, TM, JJZ

Funding acquisition, EK, TB, TR, TM, JJZ

Resources, TB, TR, TM Supervision, EK, TB, TR, TM, JJZ

Validation, EK, DP-M, LA-M, GW, RN, SB-A, MA-M, RA-S

Project administration, EK, JJZ

## Declaration of generative AI and AI-assisted technologies

During the preparation of this work, the authors used Copilot in order to proofread the manuscript. After using this tool or service, the authors reviewed and edited the content as needed and take full responsibility for the content of the publication.

## Resources table

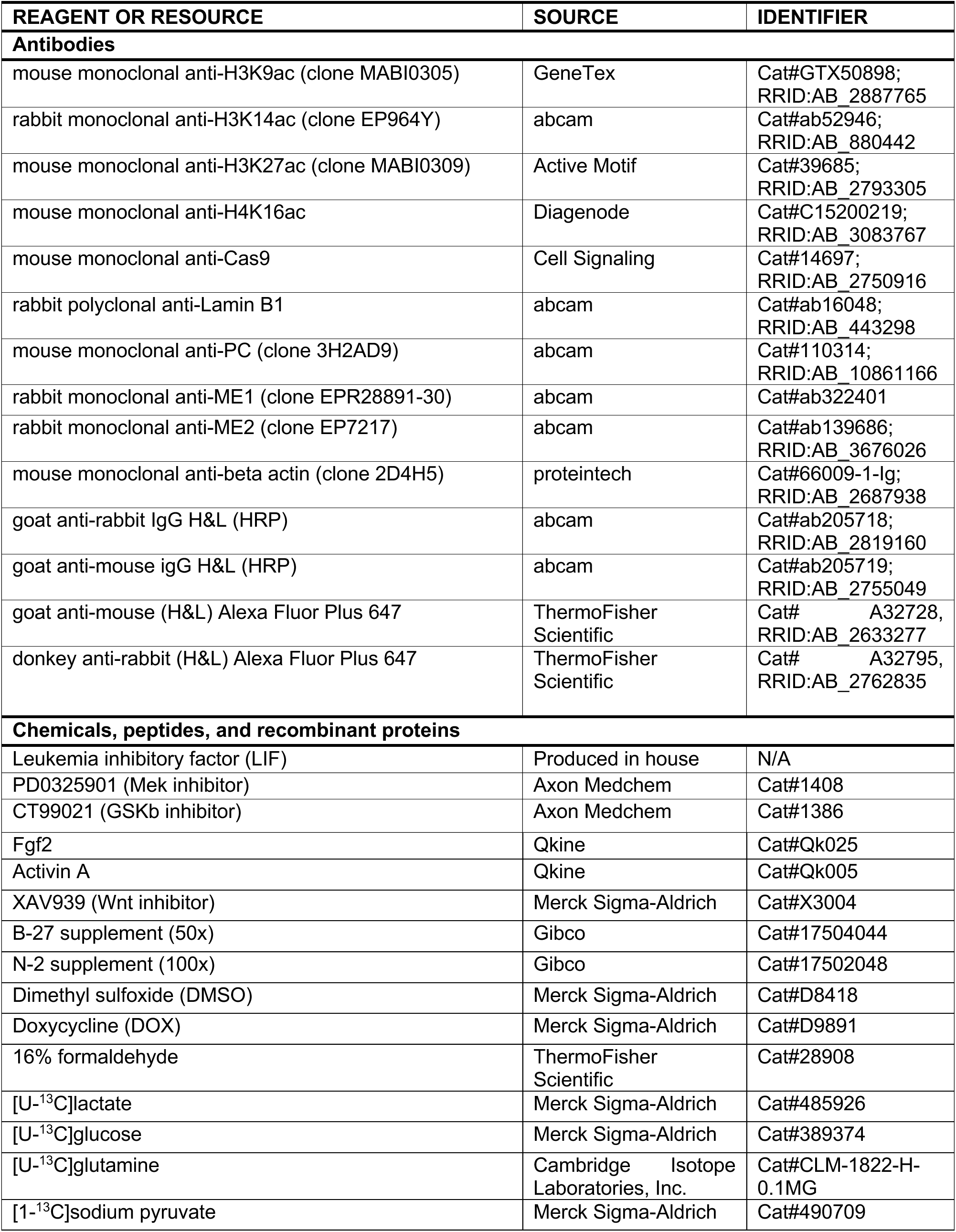
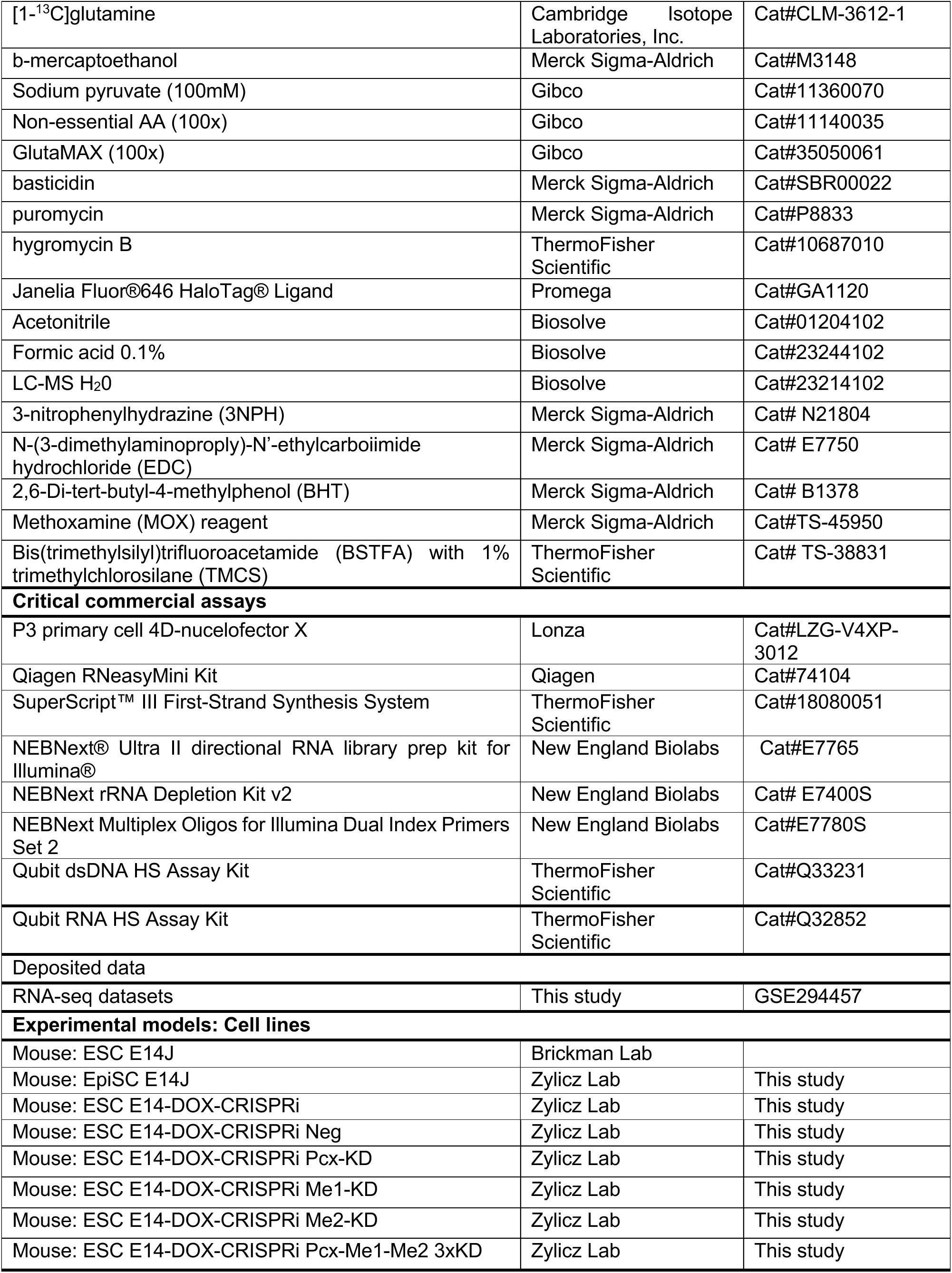
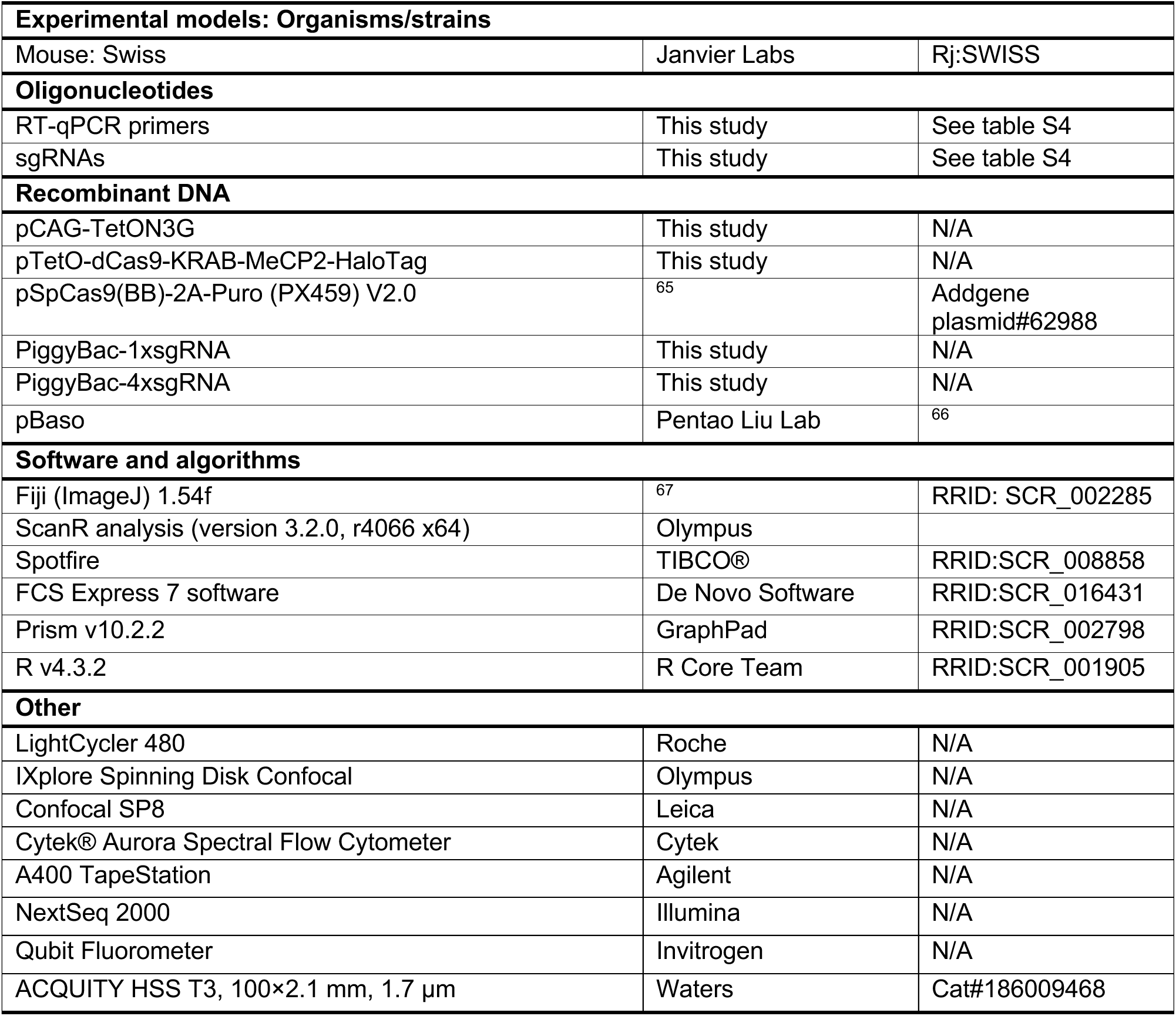

## Materials and methods

### Embryo isolation and culture

Timed natural matings were used for all experiments. Noon of the day when the vaginal plugs of mated females were identified was scored as E0.5. Outbread Rj:SWISS mice were used for all experiments. E3.25 and E6.25 embryos were collected and morphologically assessed to ensure that only viable samples were collected. Embryos were cultured for five hours at 37°C in 5% CO₂ and 5% O₂ in KSOM medium containing 95 mM NaCl, 2.5 mM KCl, 0.35 mM KH₂PO₄, 0.20 mM MgSO₄, 25 mM NaHCO₃, 1.71 mM CaCl₂, 0.01 mM EDTA, 5.5 mM glucose, 10 mM L-lactate, and 2 mM GlutaMAX. For ^13^C isotope labeling experiments, glucose, lactate or GlutaMAX were replaced with their corresponding [U-^13^C]labeled forms. Following incubation, embryos were washed with PBS supplemented with 0.01% polyvinyl alcohol (PVA), embedded in 10% gelatin, and flash frozen at -80°C until cryosectioning.

### MALDI-MSI Tissue Preparation and Matrix Deposition

Gelatin-embedded embryos were cryosectioned into 10 µm thick sections using a Cryostar NX70 cryostat (Thermo Fisher Scientific, MA, USA) at -20 °C. The sections were thaw-mounted onto indium-tin-oxide (ITO)-coated glass slides (VisionTek Systems Ltd., Chester, UK). Mounted sections were placed in a vacuum freeze-dryer for 15 minutes prior to matrix application. After drying, N-(1-naphthyl) ethylenediamine dihydrochloride (NEDC) (Sigma-Aldrich, UK) MALDI-matrix solution of 7 mg/mL in methanol/acetonitrile/deionized water (70, 25, 5 %v/v/v) was applied using a HTX M3+ Sprayer (HTX Technologies, USA). The setting was as below: temperature, 60 °C; number of passes, 20 layers; flow rate, 80 μL/min; velocity, 2000 mm/min; track spacing, 3 mm; gas flow rate 10 psi, and drying time in between passes, 30 s.

### MALDI-MSI measurements and data analysis

MALDI-TOF/TOF-MSI was performed using a RapifleX MALDI-TOF/TOF system (Bruker Daltonics GmbH, Bremen, Germany). Negative ion-mode mass spectra were acquired at a pixel size of 5 × 5 µm2 over a mass range from m/z 80-1000. Prior to analysis the instrument was externally calibrated using red phosphorus. Spectra were acquired with 30 laser shots per pixel at a laser repetition rate of 10 kHz. Data acquisition was performed using flexControl (Version 4.0, Bruker Daltonics) and visualizations were obtained from flexImaging 5.0 (Bruker Daltonics).

MSI data were exported and processed in SCiLS Lab 2024b pro (SCiLS, Bruker Daltonics) with baseline correction using convolution algorithm. All MALDI-TOF-MSI data were normalized to the Root Mean Square (RMS). Peak picking was performed (signal-to-noise-ratio > 3) on the average spectrum, and matrix peaks were excluded from the m/z feature list. The m/z values from MALDI-TOF were imported into the Human Metabolome Database (https://hmdb.ca/) after re-calibration in mMass and annotated for metabolites with an error < ±20 ppm. The 13C-labeled peaks were selected by comparing the spectrum of control and 13C-labeling experiments and annotated based on the presence of un-labeled metabolites and their theoretical m/z values. Peak intensities of the selected features were exported for all the measured pixels from SCiLS Lab, which were used for the following analysis. Natural isotope abundance correction was performed for metabolites using R package IsoCorrectoR34. Lab, which were used for the following analysis. Natural isotope abundance correction was performed for metabolites using R package IsoCorrectoR34.

For UMAP analysis, the datasets were transformed into a count matrix by multiplying the RMS-normalized intensities by 10 and taking the integer. This count data matrix was normalized and scaled using SCTransform to generate a 2-dimensional UMAP map using Seurat in R (version 4.0). All the datasets were integrated into one dataset after batch correction with rpca method in Seurat. The distribution of the pixels from different clusters on tissues were exported, and cell types were identified based on their morphology.

The average values of the exported metabolite in each cell type were calculated. The cell-type specific fraction enrichment of isotopologues was calculated based on the ratio of each 13C-labeled metabolite (isotopologue) to the sum of this metabolite abundance in each cell-type. The calculated fraction enrichment of isotopologues was used to generate pseudo-images together with pixel coordinate information exported from SCiLS Lab.

### Culture of mouse pluripotent stem cells

ESCs were maintained in naive conditions on fibronectin-coated (Corning) tissue culture plates containing N2B27 media: 1:1 ratio of DMEM/F12+Glutamax (Gibco) and Neurobasal (Gibco), 1xN2 (Gibco), 1xB27 (Gibco), 1mM Glutamax (Gibco) and 100μM b-mercaptoethanol (ThermoFisher Scientific) supplemented with 2iLIF: 3μM CT99021 (Axon Medchem), 1μM PD (Axon Medchem) and 10ng/mL LIF (made in house). For the generation of the E14-DOX-CRISPRi cell line, ESCs were cultured on gelatin-coated tissue culture plates containing Serum/LIF media based on GMEM (Sigma-Aldrich) supplemented with 10% fetal bovine serum (Sigma), 1mM Glutamax (Gibco),1x non-essential AA (Gibco), 1mM sodium pyruvate (Gibco), 100μM b-mercaptoethanol (ThermoFisher Scientific) and 10ng/mL LIF (made in house). EpiLCs and EpiSCs were cultured in FAX conditions on fibronectin-coated tissue culture plates containing N2B27 media supplemented with 20ng/mL Activin A (Qkine), 12ng/mL Fgf2 (Qkine) and 10μM XAV939 (Merck). All cell lines used were maintained under hypoxic conditions (5% O_2,_ 5% CO_2_) at 37°C and were passaged every 2-3 days.

For the ^13^C-isotope labelling metabolomic experiments, ESCs were cultured in regular N2B27 with 2iLif and EpiLCs/EpiSCs in N2B27 supplemented with FAX as described above. 5 hours prior cell harvesting, the media was replaced with N2B27 depleted of glucose, glutamine or pyruvate and supplemented with either [U-^13^C]glucose (Merck Sigma-Aldrich), [U-^13^C] glutamine(Cambridge Isotope Laboratories, Inc.), [1-^13^C] glutamine (Cambridge Istotope Laboratories, Inc.) or [1-^13^C] sodium pyruvate (Merck Sigma-Aldrich) respectively, and depending on the experiment.

### Induction of ESCs to EpiLCs

For metabolomics-related experiments, EpiLCs were generated by washing naïve ESCs three times with empty N2B27 and seeded in the appropriate density onto fibronectin-coated plates containing FAX media for 1, 2 or 3 days for the generation of EpiLCs day 1, day 2 or day 3 respectively. For experiments involving the E14-DOX-CRISPRi cell line, cells were first plated in the appropriate density and cultured in 2iLIF in the presence of DMSO or DOX (0.5μg/μl; Merck) for 24 hours. Subsequently, cells were washed three times with empty N2B27 and cultured in FAX conditions for 48 hours with DMSO or DOX. For all the experiments performed, the media was replaced every 24 hours.

### Intracellular and extracellular metabolite extractions

Prior to cell harvesting, 100 µL of spent growth medium was snap-frozen and stored at −80°C for subsequent extracellular metabolite extraction. Cells were washed three times with PBS, followed by the addition of 1.0 mL ice-cold 90% methanol. For unlabeled metabolomics, methanol was supplemented with isotopically labeled internal standards; for ^13^C isotope tracing, internal standards were omitted. Cells were collected by scraping and stored at −80°C until intracellular metabolite extraction.

For intracellular metabolite extraction, samples were thawed on ice, snap-frozen in liquid nitrogen, thawed again and vortexed. This freeze–thaw–vortex cycle was repeated three times. Samples were then incubated on ice for 1 hour and centrifuged at 15,000 rpm for 15 minutes at 4°C. For unlabeled metabolomics, 50 µL of the supernatant were transferred to GC vials and dried at room temperature using a nitrogen evaporator set at a flow of 5 L/min. For ^13^C isotope tracing, supernatants were transferred to LC vials and dried at room temperature with a nitrogen evaporator at a flow rate of 10 L/min. Dried extracts were stored at −20°C until further analysis.

For extracellular metabolite extraction, 100 µL of spent growth medium was resuspended in 200 µL of ice-cold 90% methanol containing labeled internal standards. Subsequent steps mirrored those used for intracellular metabolite extraction.

### Protein extraction and quantification

To normalize unlabeled intracellular and extracellular metabolomics data, total protein content was measured for representative samples. Briefly, cell pellets were obtained by incubating cells with TrypLE for 5 minutes, followed by the addition of growth medium and centrifugation at 1,400 rpm for 5 minutes. The resulting pellets were resuspended in RIPA buffer (50 mM Tris-HCl, pH 7.5; 150 mM NaCl; 1% NP-40; 0.1% SDS; 0.5% sodium deoxycholate; and protease inhibitor cocktail [Sigma-Aldrich, cOmplete]). Cell lysates were incubated on ice for 10 minutes, then sonicated using a Bioruptor Plus (4°C, 8 cycles of 30-second pulses with 30-second intervals). Following sonication, samples were centrifuged at 20,000 × *g* for 10 minutes at 4°C. The supernatant was collected, and protein concentration was quantified using the DC Protein Assay (Bio-Rad, 5000111).

### Gas chromatography coupled to mass spectrometry (GC-MS)

GC-MS analysis was performed on a Leco Pegasus BT GC/time-of-flight MS (St. Joseph, MI, United States) coupled to a Gerstel MPS multisampler (Mülheim an der Ruhr, Germany). A two-step derivatization reaction was performed prior to the analysis. In the first step of the reaction-methoximation, 12.5 µL of MOX reagent were added to the dried extracts, and samples were then incubated at 45°C for 1 hour. In the second step of the reaction-trimethylsilylation, methoximated samples were re-incubated with 12.5 µL of BSTFA with 1% TMCS at 45°C for 1 hour. 50 µL of 10 mg/L of 4,4’-dibromooctafluorobiphenyl in hexane were added to the samples to control for precision during injection. A Restek Rxi 5ms column (Bellefonte, PA, United States) with an inlet temperature of 270°C and a helium flow rate of 1.2 mL/min was employed for trimethylsilyl derivatives separation. Temperature gradient started at 40°C for one minute. After, temperature was increased at a rate of 20°C/min until reaching 340°C, where it was maintained for 3 minutes. Mass fragmentation spectra were generated by a 70-eV electron ionization source at an ionization current of 2.0 mA, and it was recorded at 10 Hz in the mass range 50 – 750 m/z. Together with the samples, a series of n-alkanes (C8 – C40) was analysed to calculate retention indexes (RI).

For data processing, raw GC-MS data was converted into centroid mode and exported as netCDF files. Compound identification and peak area extraction was performed with the in-house Swedish Metabolome Centre (SMC; www.swedishmetabolomicscentre.se) GC-MS software and an in-house library based on mass fragmentation spectra and retention indexes of specific compounds. Peak areas were normalized to labelled internal standards.

### Liquid chromatography coupled to mass spectrometry (LC-MS)

Extracted isotope labelled samples were derivatized with 3-nitrophenylhydrazine (3-NPH) and thereafter analysed with a 1290 Infinity II ultra-high-performance liquid chromatography (UHPLC) (Agilent Technologies, Waldbronn, Germany) coupled to a timsTOF Pro 2 mass spectrometer (Bruker Daltonics, Bremen, Germany). The derivatization occurred according to (Hodek *et al.,* 2023) with modifications. Briefly, 20 µL of 120 mmol L^−1^ EDC (dissolved in 6% pyridine and 50% methanol) and 20 µL of 200 mmol L^−1^ 3-NPH (dissolved in 50% methanol) were consecutively added to 20 µL of reconstituted samples in 1:1 methanol:water (v/v). The sample was incubated at room temperature (RT) (21°C) for 60 minutes, and afterwards 40 µL of 0.05 mg mL^−1^ BHT (dissolved in 30% pure methanol) were added to the samples and vortexed. 2 µL of each sample were analysed on an Acquity UPLC HSS-T3 column (100 × 2.1 mm, 1.8 µm, Waters, MA, USA) by using gradient elution of 0.1% formic acid (v/v) in water as mobile phase A and 0.1% formic acid in acetonitrile as mobile phase B. The flow rate was set at 0.35 mL min^−1^ with the following gradient: 0 min (5% B), 12 min (100% B), 13 min (100% B), 14 min (5% B), 17 min (5% B). Column and autosampler were kept at 30 °C and 4 °C, respectively. The mass spectrometer was operated in negative ionization mode. Ions were generated with a vacuum insulated probe heated-electrospray ionisation (VIP-HESI) source. Detection of the mass/charge ratio (m/z) of ions was set from 50 to 1000, and acquisition rates were set to 2 Hz, and the resolution was approximately 60,000. For mass calibration 50 µL internal calibrant of 10 mM Na-formate was injected at the beginning of each analysis. Data acquisition was performed with otofControl version 6.0 and Bruker Compass HyStar version 5.0 (Bruker Daltonics, Bremen, Germany). Data processing and calculation of isotopologue enrichment was performed with Bruker TASQ 2023b.

### Generation of E14-DOX-CRISPRi cell line

E14 ESCs grown in serum/Lif were nucleofected using the P3 Primary Cell 4D-Nucleofector X (Lonza) with the following 3 plasmids: px459 plasmid containing Cas9 and a sgRNA targeting the *Tigre* locus, the *pCAG-TetON3G* plasmid and *pTetO-dCas9-KRAB-MeCP2-HaloTag* plasmid. Cells resulting from this transfection were subjected to blasticidin (10μg/mL; Merck) and puromycin (1μg/mL; Merck) antibiotic selection for 5 days. Subsequently, colonies were picked, grown and treated with DOX (0.5μg/μl; Merck) for 24 hours. Next, cells were incubated with the HaloTag ligand Janelia Fluor®646 (Promega) and analyzed by flow cytometry to determine expression of dCas9-KRAB-MeCP2. The chosen clone was further validated by immunostaining and immunoblotting and retained a normal karyotype.

### Cloning of sgRNAs

The individual sgRNAs used for this study were cloned into a PiggyBac vector following the original protocol^68,69^. Muti-sgRNAs used for the generation of the Neg and 3xKD cell lines were cloned into a modified version of the original piggyBac vector following a published protocol^70^. The sgRNAs used for the study are listed in Table S4.

### Transfection of sgRNAs into E14-DOX-CRISPRi cells

ESCs were seeded onto 12-well tissue culture plates. The next day, mix A (75μl OptiMem (ThermoFisher Scientific) + 5μl lipofectamine 2000 (ThermoFisher Scientific)) and mix B (75μl OptiMem + 1μg of PiggyBac vector harboring the sgRNA(s) of interest +0.5μg of pBASO) were prepared and incubated separately for 5min at RT. Thereafter, mix A and B were combined and incubated for 15min at RT. The transfection mix was added onto the cells directly drop-by-drop. After 8 hours the media was refreshed and selection with hygromycin (160μg/mL; ThermoFisher Scientific) was started.

### RNA extraction and RT-qPCR

Total RNA was extracted using Qiagen RNeasy Mini Kit (Qiagen) with a 15min treatment of RNAse-free DNAse (Qiagen) at RT. cDNA was synthesized using Superscript III Reverse Transcriptase (ThermoFisher Scientific) following the manufacturers protocol. Subsequently, RT-qPCR was performed using PowerUp™ SYBR™ Green Master Mix (Thermo Fisher Scientific) in a LightCycler480 (Roche). Primers used for the analysis are listed in Table S4.

### Immunoblotting

For the obtention of whole cell lysates, cells were harvested and lysed in RIPA buffer (10mM Tris pH7.4, 100mM NaCl, 1mM EDTA, 1mM EGTA, 1mM NaF, 20mM Na4P2O7, 2mM Na3VO4, 1% Triton X-100, 10% glycerol, 0.1%SDS and 0.5% deoxycholate) supplemented with proteinase inhibitor cocktail tablet (Roche) and phosphatase inhibitors (Roche). Lysates were sonicated for 8 cycles (30s ON, 30s OFF). Protein quantification was performed with the Pierce^TM^ BCA protein assay kit (ThermoFisher Scientific). Protein samples were boiled in NuPAGE™ LDS Sample Buffer 4x (ThermoFisher Scientific) with 10mM dTT (ThermoFisher Scientific) at 70°C and separated in a 4-12% NuPage Bis-Tris gel (ThermoFisher Scientific). Proteins were transferred into a nitrocellulose membrane (Merck), which was blocked with 5% skimmed milk in PBST (PBS with 0.1% Tween-20) for at least 1h at RT. Subsequently, the membranes were incubated with the primary antibody overnight at 4°C. On the following day, membranes were washed three times with PBST for 10min and incubated with the corresponding HRP-conjugated secondary antibodies for 1h at RT. Thereafter, membranes were washed and developed using ECL^TM^Select western blotting detection reagent (Merck) using ChemiDoc MP Imaging System (BioRad). The intensity of the bands was quantified by ImageJ 1.54f.

### Immunostaining

Cells were cultured in 96-well imaging plates (PerkinElmer), washed with PBS and fixed with 4%formaldehyde (ThermoFisher Scientific) for 15min at RT. Subsequently, cells were washed twice with PBS, permeabilized with permeabilization buffer (0.5% Triton X-100 in PBS) for 15min at RT and blocked with blocking buffer (10% goat serum, 1% bovine serum albumin, 0.1%Tween in PBS) overnight at 4°C. Primary antibodies were diluted in blocking buffer and incubated overnight at 4°C. Next day, cells were washed three times with PBST (0.1% Tween20 in PBS) for 10min and incubated with secondary antibodies diluted in blocking buffer overnight at 4°C. Thereafter, cells were washed once with PBST and stained with DAPI (0.5μg/mL, Sigma-Aldrich) for 15min at RT. Finally, cells were washed three times with PBS before proceeding to microscope imaging and downstream analysis.

### Janelia Fluor staining and Flow cytometry

The HaloTag ligand Janelia fluor^®^ 646 (200nM, Promega) was added to the cell culture media and cells were incubated for 1h under regular cell culture conditions. Thereafter, cells were washed two times with PBS, incubated in clean media for 15 min and harvested for flow cytometry analysis. Cytek^®^ Aurora Spectral Flow Cytometer was used for flow cytometry and the data was processed and analyzed using the FCS Express 7 software (De Novo Software).

### Karyotyping

Cells were plated onto gelatin-coated 10cm dishes and treated with colcemid (0.1μg/mL, ThermoFisher Scientific) for 3 hours. Subsequently, cells were gently harvested and pelleted at 300g for 3 min and resuspended in 100μl of PBS. Next, 10mL of pre-warmed 75mM KCl was added drop by drop by pipetting to the wall of the tube and incubated for 15 min at 37°C. Thereafter, cells were spun down for 5 min at 300g using a faced angle rotor. The buffer was removed and freshly prepared fixation buffer (3:1 absolute methanol to glacial acetic acid) was added using a glass Pasteur pipette drop by drop while gently vortexing the tube. Cells were incubated at 4°C overnight. The next day, the fixation buffer was removed, leaving behind around 500mL that was used to resuspend the fixed cells. To prepare the metaphase spreads, 100μl was dropped onto a washed slide from around 20cm by tapping the pipette with force. Slides were dried for at least 30min and subsequently stained with 5% Giemsa solution (ThermoFisher Scientific) for 30 min at RT. Finally, the slides were rinsed twice with deionized water and left to dry for at least 30 min at RT. Samples were mount with 100μl of DPX mounting media (Merck).

### Bulk RNA-seq

E14-DOX-CRISPRi (Neg and 3xKD) cells were seeded in the appropriate density onto 6-well tissue culture fibronectin-coated plates in N2B27+2iLif with DMSO or DOX for 24 hours. ESCs were maintained for 48 hours more in the same conditions. For the generation of EpiLCs day 2, cells were washed three times with empty N2B27 and subsequently were cultured in FAX conditions with DMSO or DOX for 48 hours. The media was exchanged every 24 hours for all the cell lines and conditions. Puromycin (1μg/mL; Merck), blasticidin (10μg/mL; Merck) and hygromycin (160μg/mL; ThermoFisher Scientific) selection was kept throughout the length of the experiment. All cells were harvested in parallel for downstream processing. RNA was purified using the RNeasy Mini Kit (Qiagen) with a 15min treatment of RNase-Free DNase (Qiagen) at RT. 500 ng of RNA per sample was used for library preparation with the NEBNext rRNA Depletion Kit v2 (New England Biolabs), NEBNext Ultra II Directional RNA Library Prep Kit for Illumina (New England Biolabs) and NEBNext Multiplex Oligos for Illumina Dual Index Primers Set 2 (New England Biolabs). The quality of the RNA and cDNA was determined using a TapeStation (Agilent). Barcoded libraries were pooled in equimolar amounts and subjected to single-end sequencing on the Illumina NextSeq 2000 sequencer. Raw sequencing data were demultiplexed and converted into FASTQ files using Illumina bcl2fastq Conversion Software (v2.20.0.422). Resulting FASTQ files were processed using nf-core/rnaseq v3.14.0 of the nf-core collection of workflows^71^. In brief, adapters and low-quality reads were removed using Trim Galore (v0.6.7), trimmed reads were mapped to the mouse reference genome (GRCm39, Ensembl release 111) using STAR (v2.7.9a) and transcript expression was quantified with Salmon (v1.10.1)^72,73^. Transcript-level abundance estimates were imported into R (v4.3.2) and summarized to gene level using tximport package (v1.30.0)^74^. A minimal pre-filtering step was applied to retain genes with at least 10 total reads.

Differential gene expression analysis was conducted using DESeq2 (v1.28.0)^75^. Genes were considered significantly differentially expressed if they met the following criteria: an adjusted *p*-value < 0.01 and an (absolute(log_2_(fold change)) > 0.58 across the specified comparisons. Gene set enrichment analysis for the significantly upregulated and downregulated genes in the 3xKD versus Neg in EpiLCs was performed using the clusterProfiler (v4.10.1) package and visualized with the GseaVis (v0.0.5) package^76,77^.

For *K*-means clustering, the likelihood ratio test (LRT) from the DESeq2 package was used to identify differentially expressed genes, with significance defined as an adjusted *p*-value < 0.01. *K*-means clustering was performed on the resulting matrix of rlog transformed counts using the pheatmap package (v1.0.12). The same package was utilized to visualize the results as heatmap. Gene Ontology (GO) and KEGG pathway enrichment analyses were performed for each gene cluster using the clusterProfiler package and/or Cytoscape^78^. The analysis focused on the Biological Process ontology and KEGG pathways, with significance defined as an adjusted *p*-value < 0.01.

### Colony formation assay and alkaline phosphatase staining

EpiLCs day 2 were generated as described above. Cells were washed thoroughly with N2B27 three times and seeded at low density in naïve culture conditions. Five days later, colonies were fixed and stained with leukocyte alkaline phosphatase kit (Merck) following the manufacturer’s protocol. AP+ colonies were imaged using GelDoc XR^+^(BioRad), identified and quantified by ImageJ 1.54f.

### Histone enrichment and quantification of histone PTMs by MS

ESCs, EpiLCs day 2 and EpiSCs were cultured for 48 hours under the corresponding media conditions and as described above. 5 hours prior harvesting, the media was replaced with fresh regular media (for the unlabeled samples) or with N2B27 depleted of glucose or glutamine and supplemented with [U-13C]glucose (Merck Sigma-Aldrich) or [U-13C]glutamine (Cambridge Isotope Laboratories, Inc.) respectively. Thereafter, cells were resuspended vigorously in 1ml of nuclei isolation buffer (0.5mM PMSF, 5µg/ml Leupeptin (Merck Sigma-Aldrich), 5µg/ml Aprotinin (Merck Sigma-Aldrich), 5mM Na-butyrate, 0.1% Triton) to facilitate rupture of the membranes. The nuclei were isolated by centrifugation for 15 min at 2300g at 4°C. Next, the nuclear pellet was resuspended with 100μl of nuclei isolation buffer supplemented with 0.1% SDS and 250 U of benzonase (Merck Sigma-Aldrich) and incubated for 5 min at 37°C to digest nuclear extracts. Protein concentration was measured using DC Protein Assay (Bio-Rad, 5000111).

Approximately 4 mg of histone octamer were mixed with an approximately equal amount of a heavy-isotope labelled histone standard and were separated on a 17% SDS-PAGE gel^79^. Histone bands were excised, chemically acylated with propionic anhydride and in-gel digested with trypsin, followed by peptide N-terminal derivatization with phenyl isocyanate (PIC)^80^ Peptide mixtures were separated by reversed-phase chromatography on an EASY-Spray column (Thermo Fisher Scientic), 25-cm long (inner diameter 75 µm, PepMap C18, 2 µm particles), which was connected online to a Q Exactive Plus instrument (Thermo Fisher Scientific) through an EASY-Spray™ Ion Source (Thermo Fisher Scientific), as described^80^. The acquired RAW data were analyzed using Epiprofile 2.0^81^, selecting the “histone_C13” and “SILAC” options, followed by manual validation^80^. For each acetylated peptide, a % 13C relative abundance (%RA) was estimated by dividing the area under the curve (AUC) of 13C-labeled peptides for the sum of the areas corresponding to all the observed labelled/unlabeled forms of that peptide and multiplying by 100. For the experiment shown in Figure S4G, the levels of common histone acetylations and methylations were quantified as described^80^. The mass spectrometry data have been deposited to the ProteomeXchange Consortium^82^via the PRIDE partner repository with the dataset identifier PXD062776.

### Image analysis and quantification

Images from the immunostainings were acquired using an Olympus IXplore Spinning Disk Confocal microscope using an Olympus UPLXAPO 20×0.80– WD 0.60mm objective. For each well in a 96-well plate, 12 images composed of 30 Z-fields separated by 1μm each were taken. The ScanR analysis software (version 3.2.0, r4066 x64, Olympus) was used for image segmentation and quantification of the following parameters: X, Y, area, well, circularity, median, average and total fluorescent intensities of the corresponding antibodies. Subsequently, the data was imported to the program TIBCO^®^ Spotfire for further analysis.

### Statistical analysis

All the experiments were performed at least in biological triplicates, unless otherwise stated. The statistical tests were performed using R and GraphPad Prism 10. The statistical details for each experiment can be found in the figure legends.

