## Supplemental Information for "TCA cycle rewiring underpins implantation and histone acetylation programming"

### Supplementary figures

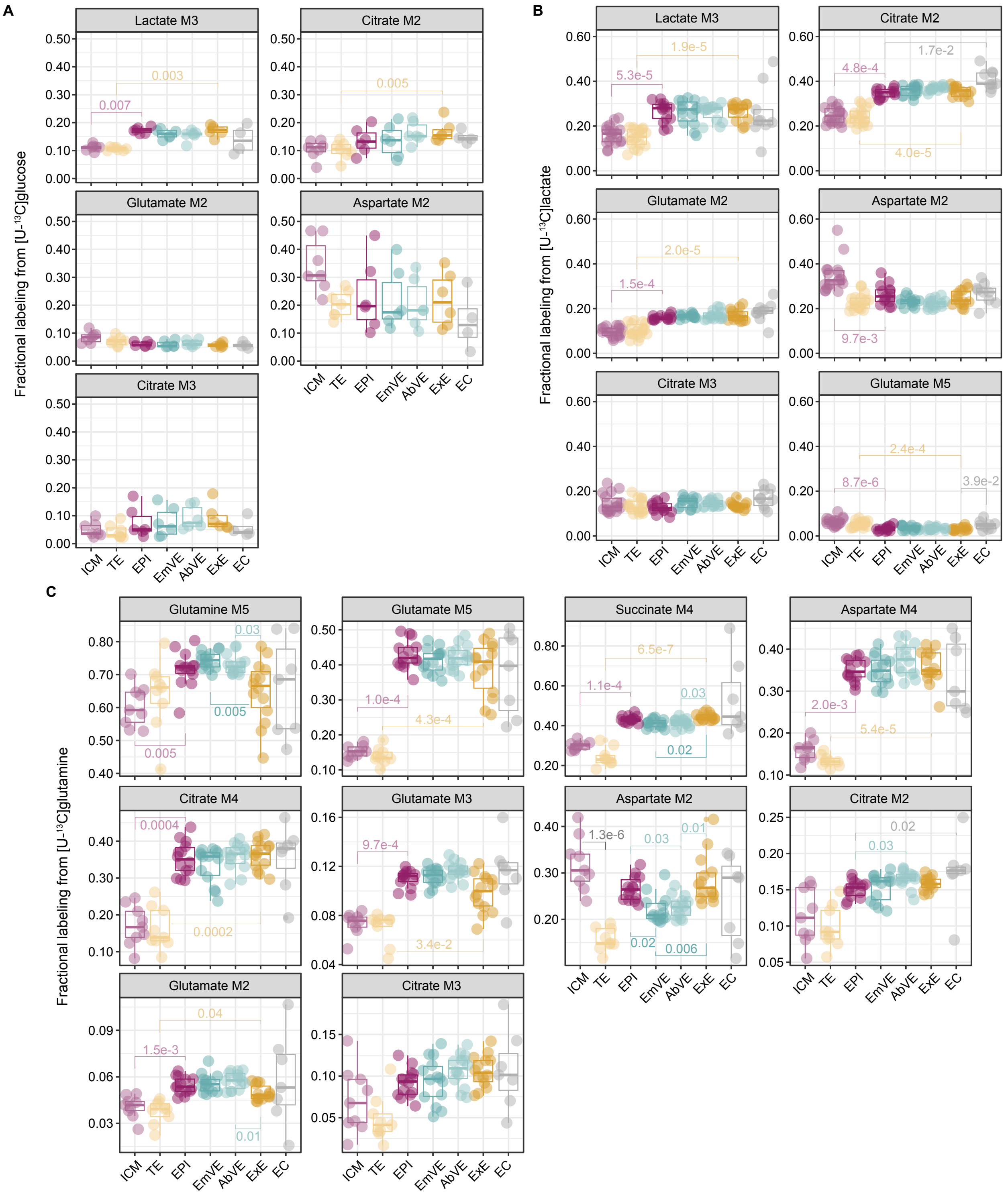


### Supplementary Figure 1

(**A-C**) Fractional labeling of metabolites derived from [U-$^{13}$C]glucose (A), [U-$^{13}$C]lactate (B) or [U-$^{13}$C]glutamine (C) in spatially resolved lineages in E3.5 and E6.5 embryos. (A) ICM, n = 7; TE, n = 7; EPI, n = 6; EmVE, n = 6; AbVE, n = 6; ExE, n = 6; EC, n = 4. (B) ICM, n = 18; TE, n = 18; EPI, n = 14; EmVE, n = 14; AbVE, n = 14; ExE, n = 14; EC, n = 9. (C) ICM, n = 9; TE, n = 9; EPI, n = 14; EmVE, n = 14; AbVE, n = 14; ExE, n = 14; EC, n = 7. ICM, inner cell mass; TE, trophectoderm; EPI, epiblast; EmVE, embryonic visceral endoderm; AbVE, abembryonic visceral endoderm; ExE, extraembryonic ectoderm; EC, ectoplacental cone. Data points correspond to individual embryos. Statistical significance was assessed using the Kruskal-Wallis test followed by Dunn's post-hoc test, with p-values adjusted with the Benjamini-Hochberg (BH) correction method.

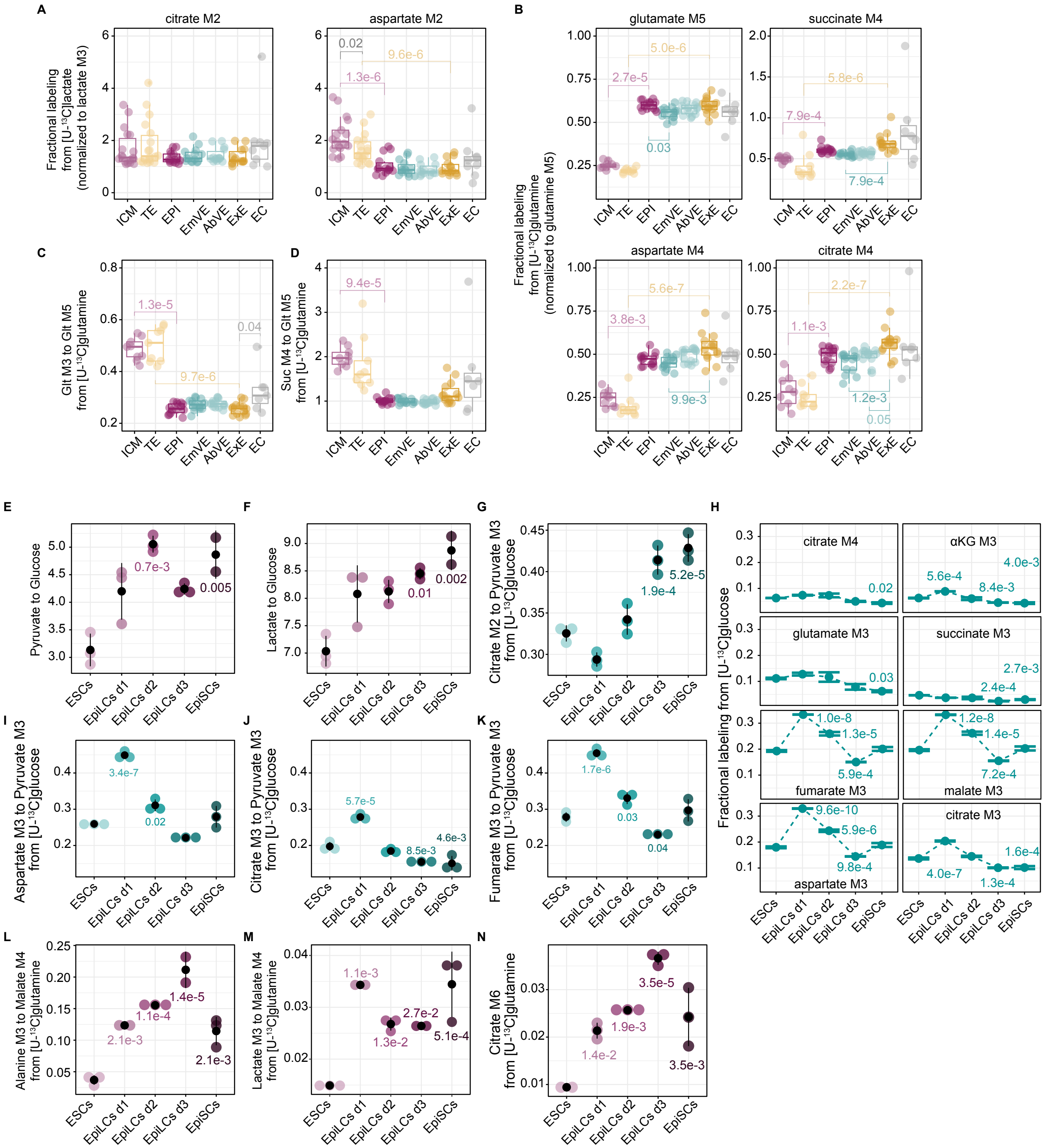


**Supplementary Figure 2**

(**A-B**) Fractional labeling of metabolites derived from [U-$^{13}$C]lactate (A) and [U-$^{13}$C]glutamine (B) in spatially resolved lineages in E3.5 and E6.5 embryos. The values are normalized to M3 lactate (A) and M5 glutamine (B). (A) ICM, n = 18; TE, n = 18; EPI, n = 14; EmVE, n = 14; AbVE, n = 14; ExE, n = 14; EC, n = 9. (B) ICM, n = 9; TE, n = 9; EPI, n = 14; EmVE, n = 14; AbVE, n = 14; ExE, n = 14; EC, n = 7. (**C**) Relative oxidative TCA activity in spatially resolved lineages in E3.5 and E6.5 embryos, represented by the fractional labeling of M3 glutamate normalized to M5 glutamate from [U-$^{13}$C]glutamine. ICM, n = 9; TE, n = 9; EPI, n = 14; EmVE, n = 14; AbVE, n = 14; ExE, n = 14; EC, n = 7. (**D**) Fractional labeling of M4 succinate normalized to M5 glutamate from [U-$^{13}$C]glutamine in spatially resolved lineages in E3.5 and E6.5 embryos. ICM, n = 9; TE, n = 9; EPI, n = 14; EmVE, n = 14; AbVE, n = 14; ExE, n = 14; EC, n = 7. (**E-F**) Ratio of pyruvate (E) and lactate (F) to glucose based on intracellular metabolite abundances throughout pluripotency progression (n = 3). (**G**) Relative glucose entry to the TCA cycle through the activity of pyruvate dehydrogenase throughout pluripotency progression, represented by the ratio of M2 citrate to M3 pyruvate from [U-$^{13}$C]glucose (n = 3). (**H**) Fractional labelling of TCA cycle intermediates from [U-$^{13}$C]glucose throughout pluripotency progression, focusing on isotopologues derived from the second turn of the cycle (n = 3). (**I-K**) Relative pyruvate carboxylase activity throughout pluripotency progression, represented by the fractional labeling of M3 isotopologues of aspartate (I), citrate (J) and fumarate (K) normalized to M3 pyruvate from [U-$^{13}$C]glucose (n = 3). (**L-M**) Relative malic enzyme activity throughout pluripotency, represented by the fractional labeling of M3 isotopologues of alanine (L) and lactate (M) normalized to M4 malate from [U-$^{13}$C]glutamine (n = 3). (**N**) Fractional labeling of M6 citrate from [U-$^{13}$C]glutamine throughout pluripotency (n = 3). (**A-D**) ICM, inner cell mass; TE, trophectoderm; EPI, epiblast; EmVE, embryonic visceral endoderm; AbVE, abembryonic visceral endoderm; ExE, extraembryonic ectoderm; EC, ectoplacental cone. Data points correspond to individual embryos. Statistical significance was assessed using the Kruskal-Wallis test followed by Dunn's post-hoc test, with p-values adjusted with the Benjamini-Hochberg (BH) correction method. (**E-N**) Data represent the mean of three biological replicates ± standard deviation (SD) (E-G, I-N) or ± standard error of the mean (SEM) (H) per condition, with each colored point indicating an individual biological replicate. Statistical significance was assessed using one-way ANOVA followed by Tukey's HSD post-hoc test. Levene's test was used to evaluate homogeneity of variances, and Shapiro-Wilk test was applied to assess the normality of residuals for each metabolite.

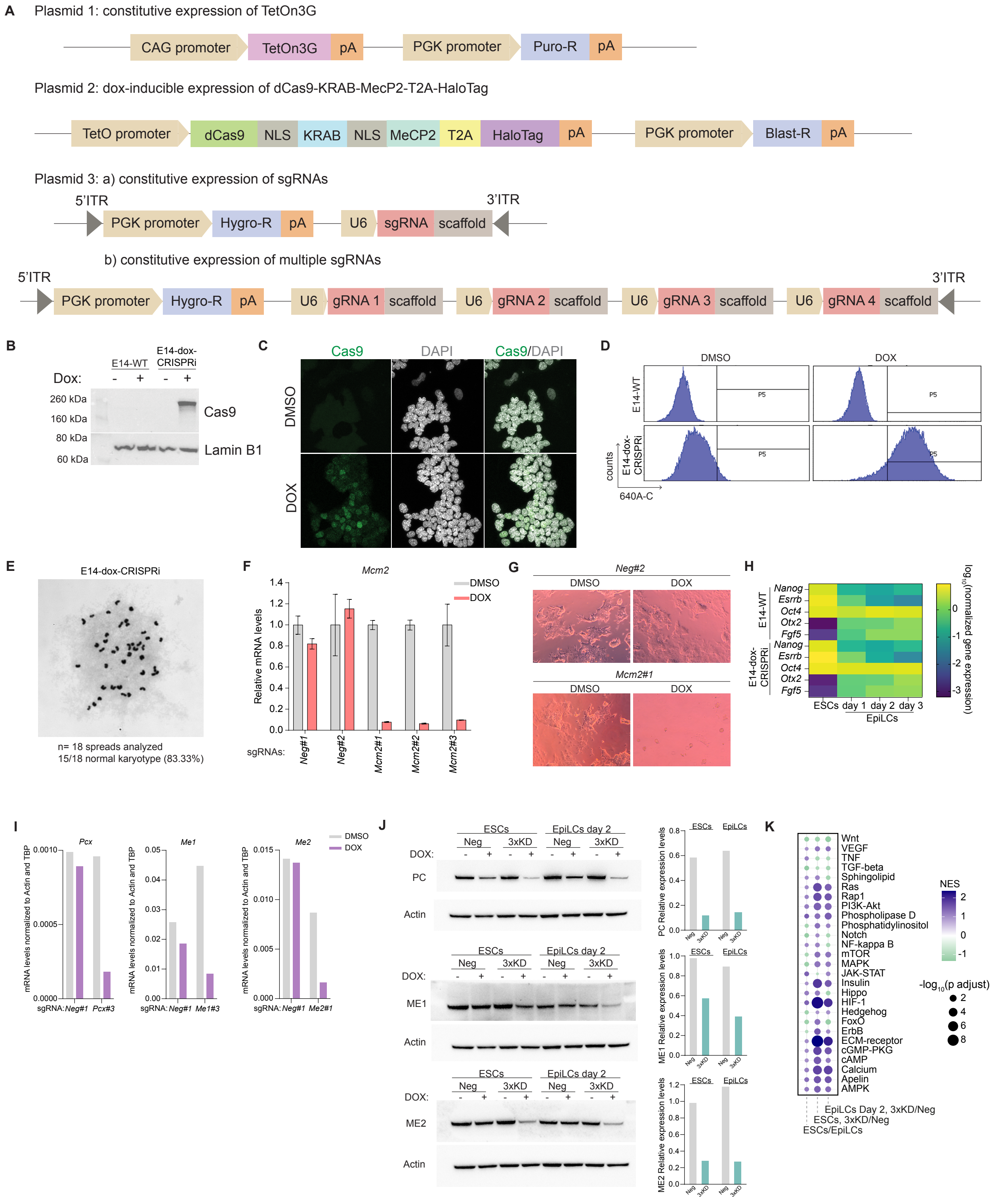


### Supplementary Figure 3

(**A**) Plasmids employed for the generation of the E14-DOX-CRISPRi cell line. Plasmid 1: Integrated into one *Tigre* allele for constitutive expression of the TetOn3G DOXycycline(DOX)-binding transactivator protein under the control of the CAG promoter. This plasmid also includes a puromycin resistance cassette driven by the PGK promoter. Plasmid 2: Integrated into the other *Tigre* allele. Upon DOX treatment, TetOn3G binds to the TetO promoter, inducing transcription of the dCas9-KRAB-MeCP2-T2A-HaloTag. The T2A self-cleaving peptide facilitates the separation of dCas9-KRAB-MeCP2 and HaloTag proteins after translation. Additionally, this plasmid contains a blasticidin resistance cassette driven by the PGK promoter. Plasmid 3: Represents the piggyBAC plasmids, with plasmid 3a encoding a single sgRNA, and plasmid 3b accommodating up to four sgRNAs. Constitutive expression of sgRNA(s) guides the dCas9-KRAB-MeCP2 complex to the transcriptional start site of the target genes. Plasmid 3 also harbors a hygromycin resistance cassette under the PGK promoter. (**B**) Immunoblot analysis of whole-cell lysates from E14-WT and E14-DOX-CRISPRi ESCs following 24-hour treatment with either DMSO or DOX. Samples were probed with Cas9 and Lamin B1 antibodies. (**C**) Immunofluorescence analysis of E14-DOX-CRISPRi ESCs after 24 hours of treatment with either DMSO or DOX. Cells were stained with a Cas9 antibody (green). (**D**) Flow cytometry histogram analysis of Janelia Fluor 646 (HaloTag ligand) in E14-WT and E14-DOX-CRISPRi ESCs after 24 hours of treatment with DMSO or DOX. (**E**) Bright field image of a chromosome spread used for karyotyping the E14-DOX-CRISPRi cell line. Analysis of 18 metaphase spreads revealed that 83.33% exhibited a normal karyotype of 40 chromosomes. (**F**) RT-qPCR analysis of *Mcm2* expression in the indicated cell lines following 72 hours of DMSO (gray) or DOX (red) treatment. mRNA levels were normalized to the expression of the housekeeping genes *actin* and *tbp*. (**G**) Representative bright-field images of E14-DOX-CRISPRi cells transfected with either *Neg#2* or *Mcm2#1* sgRNAs following 5 days of DMSO or DOX treatment. (**H**) Heatmap displaying log10-normalized gene expression of key pluripotency genes in E14-WT and E14-DOX-CRISPRi ESCs and EpiLCs (day 1 to day 3). (**I**) RT-qPCR analysis of the indicated genes in the specified cell lines following 72 hours of DMSO (gray) or DOX (purple) treatment. mRNA levels were normalized to the expression of housekeeping genes *actin* and *tbp*. (**J**) Immunoblot analysis of whole-cell lysates from E14-DOX-CRISPRi (Neg and 3xKD) ESCs and EpiLCs (day 2) following 72 hours of DMSO or DOX treatment. Blots were probed with antibodies for PC, ME1, ME2 and actin. The intensity of the bands was measured using ImageJ 1.54f. The levels of PC, ME1 and ME2 were normalized to the respective actin loading control. Bar plots show the relative expression levels of DOX-treated samples compared to their respective DMSO counterparts. (**K**) Gene set enrichment analysis of significantly differentially expressed genes, focusing on signaling pathways from the KEGG database. Comparisons include Neg ESCs versus Neg EpiLCs (column 1), 3xKD versus Neg in ESCs (column 2), and 3xKD versus Neg in EpiLCs day 2 (column 3). The normalized enrichment score (NES) is represented by a color gradient, while the $-\log_{10}$transformed adjusted *p* values are indicated by circle size.

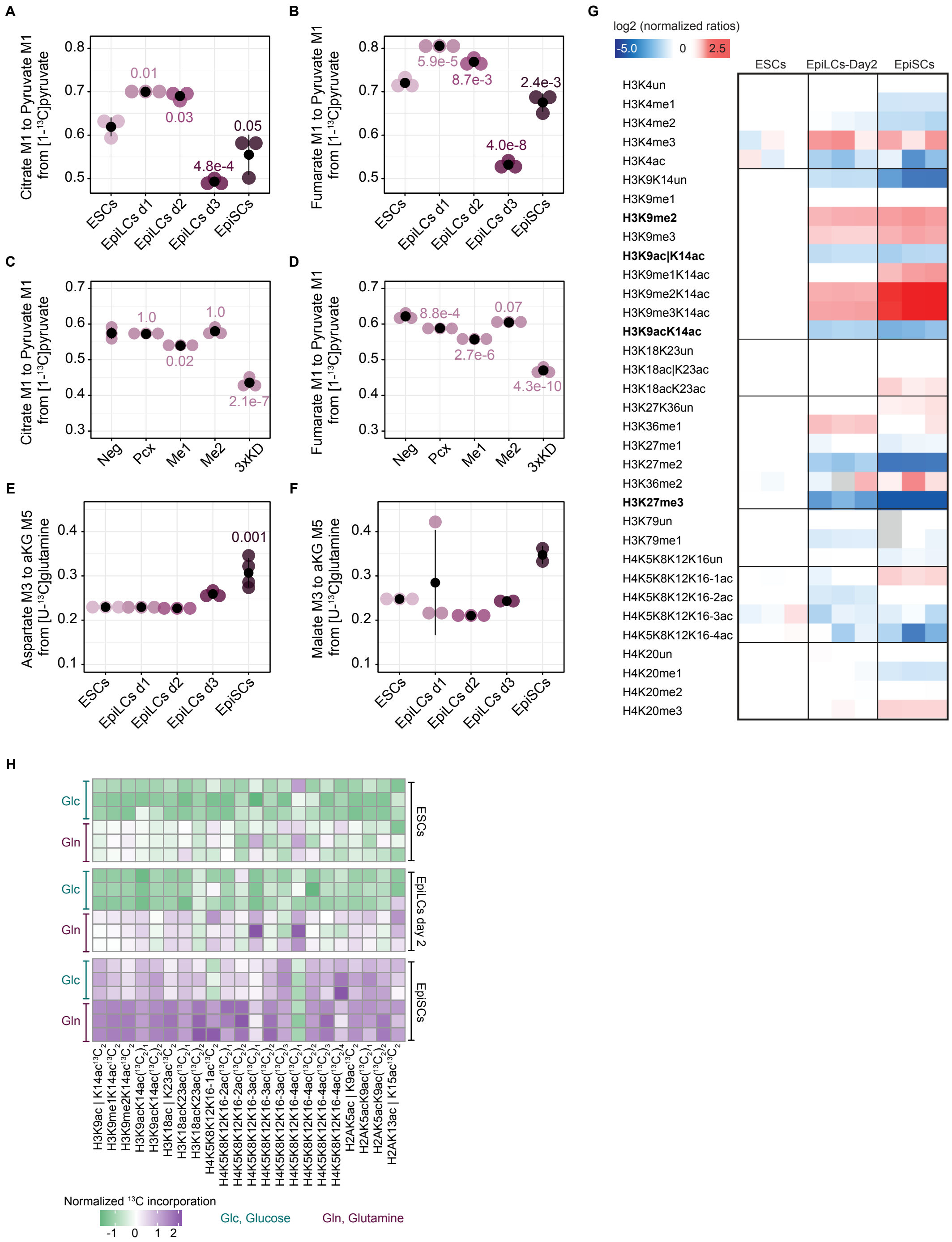


### Supplementary Figure 4

(**A-B**) Relative pyruvate carboxylase activity throughout pluripotency progression, represented by the fractional labeling of M1 isotopologues of citrate (A) and fumarate (B) normalized to M1 pyruvate from [1-$^{13}$C]pyruvate (n = 3). (**C-D**) Relative pyruvate carboxylase activity in Neg EpiLCs day 1 and following the repression of Pcx, Me1, Me2, and all three genes (3xKD). Activity is represented by the fractional labeling of M1 isotopologuea of citrate (C) and fumarate (D) normalized to M1 pyruvate from [1-$^{13}$C]pyruvate (n =

3). (**E-F**) Fractional labeling of M3 isotopologues of aspartate (E) and malate (F) normalized to M5 aKG from [U-$^{13}$C]glutamine throughout pluripotency progression (n = 3). (**G**) Heatmap showing L/H ratios (where L=sample and H=internal standard) for specific histone post-translational modifications across ESCs, EpiLCs day 2, and EpiSCs. The data has been normalized to the average levels of the corresponding histone peptide measured in ESCs (n = 3). (**H**) Heatmap showing the normalized $^{13}$C-incorporation for differentially modified histone peptides derived from [U-$^{13}$C]glucose (Glc) or [U-$^{13}$C]glutamine (Gln) in ESCs, EpiLCs day 2 and EpiSCs. Residues separated by "l" indicate the measurement of one residue or the other (n = 3). (**A-F**) Data represent the mean of three biological replicates ± standard deviation (SD) per condition, with each colored point indicating an individual biological replicate. Statistical significance was assessed using one-way ANOVA followed by Tukey's HSD post-hoc test. Levene's test was used to evaluate homogeneity of variances, and Shapiro-Wilk test was applied to assess the normality of residuals for each metabolite.

**Supplementary tables**

**Table S1**

$^{13}$C metabolite labeling and metabolite abundances in embryos, stem cells and following the repression of the genes *Pcx*, *Me1*, *Me2*and all three (3xKD) in ESCs and EpiLCs. Related to Figures 1, 2, 3A, 3B, 3E, 3F, 4, 5A and 5B.

**Table S2**

*K*-means clusters with gene ontology (GO) analysis for the transcriptional changes in 3xKD versus Neg in ESCs and EpiLCs day 2. Only the top GO terms are shown. Related to Figure 3G.

**Table S3**

DSeq2 analysis for the transcriptional changes in 3xKD versus Neg in ESCs and EpiLCs day 2. Only the statistically significant results (adjusted *p* value < 0.01 and a $\log_2$ transformed fold change > 0.58 or < -0.58) for the different comparisons are shown. Related to Figures 3H, 3I and 3K.

**Table S4**

Oligonucleotides used for this study. Related to Figure 3 and 4J.
